# Insights into the structure and modulation of human TWIK-2

**DOI:** 10.1101/2025.02.19.639014

**Authors:** Qianqian Ma, Ciria C. Hernandez, Vikas Navratna, Arvind Kumar, Abraham Lee, Shyamal Mosalaganti

## Abstract

The *<u>T</u>andem of pore domain in a <u>W</u>eak Inward <u>R</u>ectifying <u>K</u>^+^ channel 2* (TWIK-2; KCNK6) is a member of the Two-Pore Domain K*^+^* (K_2P_) channel family, which is associated with pulmonary hypertension, lung injury, and inflammation. The structure and regulatory mechanisms of TWIK-2 remain largely unknown. Here, we present the cryo-electron microscopy (cryo-EM) structure of human TWIK-2 at ∼3.7 Å and highlight its conserved and unique features. Using automated whole-cell patch clamp recordings, we demonstrate that gating in TWIK-2 is voltage-dependent and insensitive to changes in the extracellular pH. We identify key residues that influence TWIK-2 activity by employing structure and sequence-guided site-directed mutagenesis and provide insights into the possible mechanism of TWIK-2 regulation. Additionally, we demonstrate the application of high-throughput automated whole-cell patch clamp platforms to screen small molecule modulators of TWIK-2. Our work serves as a foundation for designing high-throughput small molecule screening campaigns to identify specific high-affinity TWIK-2 modulators, including promising new anti-inflammatory therapeutics.

## Introduction

Two-pore-domain (K_2P_) channels are a diverse family of potassium (K^+^) selective ion channels that supervise background K^+^ currents. They are critical to maintaining membrane resting potential and regulating cellular excitability ^1–3^. In mammals, the K_2P_family has 15 members classified into six subgroups based on their functional diversity: TWIK, TREK, TRESK, TASK, TALK, and THIK ^4,5^. Unlike Voltage-gated K^+^(K_v_) channels, K_2P_ channels lack the canonical voltage-sensing domain. While the overall architecture of K_2P_ channels is evolutionarily conserved, multiple structural variations and modulation sites make these channels susceptible to regulation by voltage-independent factors such as temperature, pH, pressure, bioactive lipids, and volatile anesthetics ^5–7^.

Among the 15 members of the K_2P_family, channels from the TREK and TASK subfamilies have been extensively structurally characterized ^8–13^, offering profound insights into their gating mechanisms, physiological functions, and pharmacological potential ^8,14–24^. TREK channels are known for their mechano- and thermosensitivity and role in sensory response ^14,18,25,26^, and TASK channels are well-characterized for their pH sensitivity and role in maintaining neuronal excitability ^12,13,27–29^. Nevertheless, the gating mechanisms, physiological modulation, and pharmacological potential of the TWIK family remain underexplored primarily because of the weak currents ^30,31^ and poor heterologous expression of these channels ^32–34^.

TWIK-1 (KCNK1) is the first and the only structurally characterized mammalian K_2P_ channel member of the TWIK family ^30,35^. The poor currents in TWIK-1 were initially believed to be due to intracellular localization resulting from SUMOylation ^36^. However, mutations in the SUMOylation site and over-expression of a recombinant plasma-membrane-localized TWIK-1 demonstrated that the poor currents are characteristic of TWIK family members ^37,38^. Recent studies involving a combination of electrophysiology, site-directed mutagenesis, molecular dynamics simulations, and structural studies of TWIK-1 at low (pH 5.5) and neutral pH (pH 7.4) revealed the molecular basis for pH-mediated regulation of gating and conformational dynamics of the selectivity filter in TWIK-1^39,40^.

TWIK-2 (KCNK6) shares ∼54% sequence similarity with TWIK-1 and exhibits ubiquitous tissue distribution ^31,41^. TWIK-2 is implicated in diverse pathological conditions such as pulmonary hypertension, acute lung injury, hearing loss, and NLRP3 inflammasome-induced inflammation ^42–46^. Several groups have reported a significant variation in the current recordings for TWIK-1 and TWIK-2, underscoring the challenges of robust electrophysiological studies for the TWIK subfamily ^31,33,41^. The weak basal activity of TWIK-2 could be due to its subcellular partitioning between the plasma membrane and endolysosomes. Additionally, it has been reported that Y308A mutation within the lysosomal-targeting YXXØ motif in TWIK-2 promotes plasma membrane expression of TWIK-2^34^. *Kcnk6* knockout studies have shown suppression of NLRP3 inflammasome activation in macrophages, implicating endolysosomal localization of TWIK-2 in inflammation ^45,46^. Despite the high sequence similarity with TWIK-1, TWIK-2 is believed to be insensitive to pH and indifferent to activation by phosphorylation ^30,31,38,40^. The mechanism of regulation of TWIK-2 is poorly understood because of the lack of three-dimensional structures and inconsistent reports of its electrophysiological recordings ^31,34,41^. Additionally, although substantial progress has been made in developing K_2P_ channel modulators, particularly for the well-characterized channels of the TREK subfamily, there is a dearth of high-affinity selective modulators of TWIK-2. The small molecule modulators of TWIK-2 activity, such as ML365 and NPBA, also cross-react with other K_2P_ channels ^5,25,47–52^

To gain deeper insights into the molecular basis of channel regulation and to aid the pharmacological characterization of TWIK-2, we determined the structure of full-length human TWIK-2 by single particle cryo-electron microscopy (cryo-EM). We uncovered conserved and unique features of this K_2P_ channel subtype. Using a combination of structure-guided site-directed mutagenesis and automated high-throughput whole-cell patch clamp electrophysiology, we identified key amino acids that differentiate TWIK-2 from TWIK-1 and other members of the K_2P_ family. Finally, using the existing K_2P_ modulators, we developed an effective strategy for high-throughput screening of TWIK-2 modulators.

## Results

### Heterologous expression of recombinant human TWIK-2

We expressed full-length TWIK-2 (hereafter TWIK-2) in HEK293 GnTI^-^ cells and verified that it was functional by whole-cell patch-clamp electrophysiology (Figure 1A and S2A). When recorded in high potassium (100 mM) at pH 7.4, the TWIK-2 channels evoked significantly larger outward potassium currents than the non-transfected (non-T) cells (Figure 1A, p < 0.0001). TWIK-1 is sensitive to pH ^39,40^. To assess if TWIK-2 behaves similarly, we tested the conductance at two different pHs (7.4 and 5.5) in high potassium (Figure 1B). We did not observe significant changes in peak current amplitudes in TWIK-2 at pH 7.4 and pH 5.5 in these conditions (p = 0.620), demonstrating that TWIK-2 is insensitive to changes in extracellular pH. Notably, TWIK-2 channels elicited outward potassium currents that increased with depolarization (over +30 mV), suggesting voltage-dependent activation of TWIK-2 (Figure 1A and 1B).

**Figure 1.**
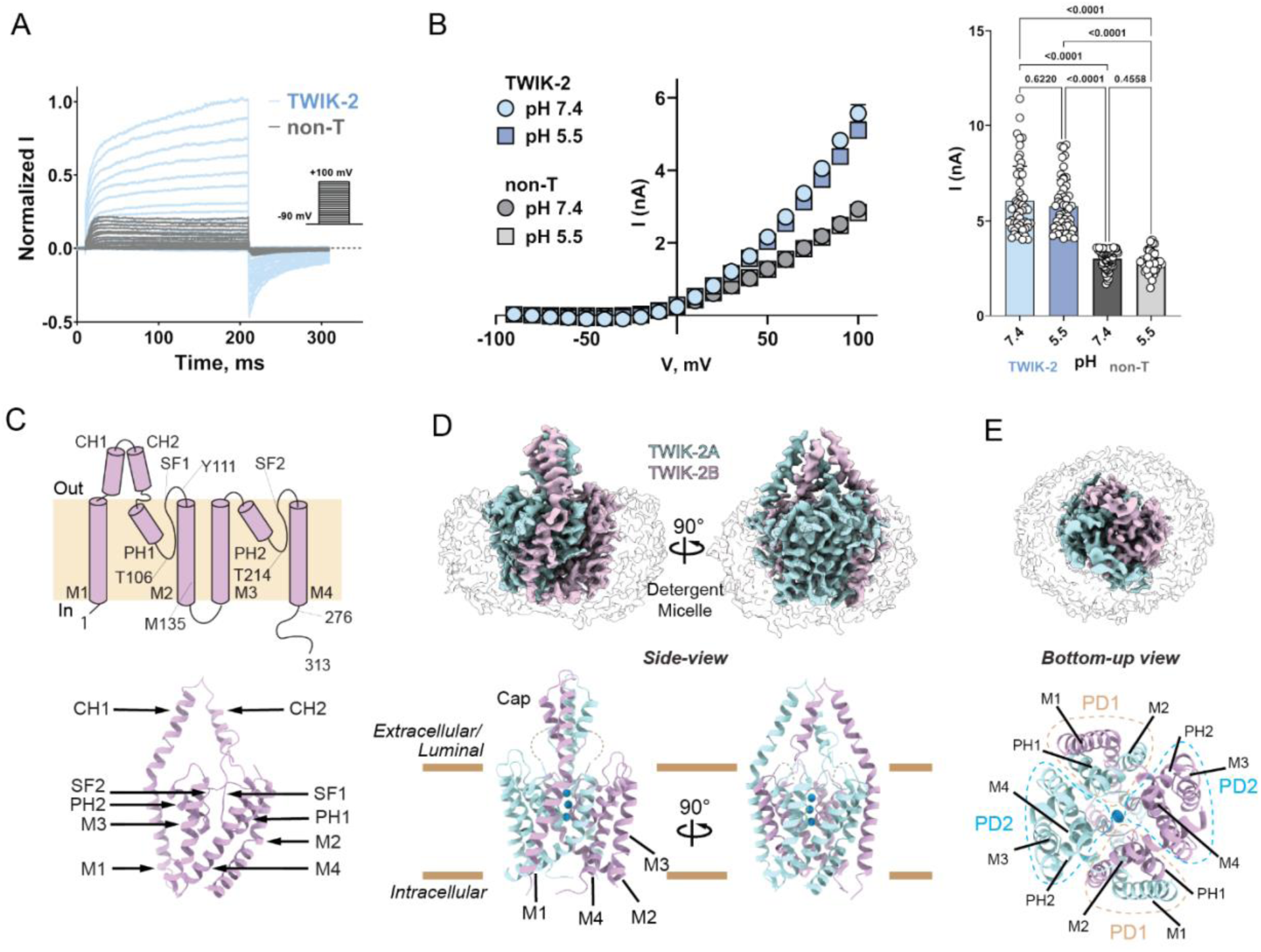
Function and structure of TWIK-2. **A.** The normalized current (I) traces as a function of time (ms). 200-ms voltage steps were performed ranging from −90 to +100 mV, starting from a holding potential of −90 mV using whole-cell patch clamp configuration recorded for non-transfected cells (non-T) and cells expressing TWIK-2 under high extracellular potassium (100 mM K^+^) at pH 7.4. **B.** Current-voltage (I/V) plots for the maximum peak currents of cells overexpressing TWIK-2 or non-T cells under high extracellular potassium at pH 7.4 and 5.5. The data are presented as mean ± SEM for TWIK-2 (pH 7.4, n = 65; pH 5.5, n = 62) and non-T (pH 7.4, n = 32; pH 5.5, n = 39). Scatter plots illustrate the peak currents obtained at +100 mV for TWIK-2 (pH 7.4, n = 52; pH 5.5, n = 52) and non-T (pH 7.4, n = 60; pH 5.5, n = 39). The data are shown as mean ± SD. Statistical analysis was performed using a two-way ANOVA with multiple comparisons. **C.** Topology and model of TWIK-2 protomer showing transmembrane helices (M1-M4), cap-forming helices (CH1 and CH2), pore-forming helices (PH1 and PH2), selectivity filters (SF1 and SF2). Critical residues along the ion permeation pathway are marked. **D**. Cryo-EM map and model of TWIK-2 dimer. The protomers, TWIK-2 A and B, are colored light magenta and light cyan, respectively. DMNG micelle density is rendered transparent. Cartoon representation of TWIK-2 in an *en-face* view parallel to the membrane. K^+^ ions are highlighted as blue spheres. **E**. Intracellular view of cryo-EM map (top) and model of TWIK-2 (bottom) with pore domains (PD1: orange and PD2: blue). Note M1s in PD1s are domain-swapped.

K_2P_ channels show poor sequence conservation and high variability in the length of their C-terminal regions (Figure S1). Removing the unstructured C-terminal region has been predicted to increase protein expression and stability of other K_2P_ channels ^12,27^. To assess the role of the extended C-terminus in TWIK-2, we deleted the terminal 46 residues and generated an ΔC-TWIK-2 construct (1-276 amino acids). We compared the channel currents and stability of ΔC-TWIK-2 with TWIK-2 using automated whole-cell patch clamp assay and fluorescence-detection size-exclusion chromatography (FSEC) (Figure S2). There was no significant change in the current density of ΔC-TWIK-2 compared to TWIK-2, suggesting a similar expression level and channel activity (Figure S2A-S2D). However, upon solubilization in 1% detergent followed by ultracentrifugation and analysis by FSEC, we noticed a poor yield for the ΔC-TWIK-2 construct compared to full-length TWIK-2, suggesting low stability in detergent micelles (Figure S2F). Based on our whole-cell recordings and FSEC analysis, we proceeded with the full-length human TWIK-2 construct for overexpression and subsequent structural studies. We purified TWIK-2 using Decyl maltose neopentyl glycol (DMNG) at pH 7.5 and 150 mM KCl and confirmed the protein homogeneity and purity by size-exclusion chromatography and SDS-PAGE analysis (Figure S3A). The peak fractions, corresponding to TWIK-2 dimer, that eluted at ∼12.4 ml on a Superdex 200 Increase size-exclusion chromatography column were used for subsequent structural analysis (Figure S3B).

### Molecular architecture of TWIK-2

We determined the structure of TWIK-2 at ∼3.7 Å (Supplementary Table S1 and Figure S3C-S3F). We were able to reliably model all the secondary structure elements of TWIK-2 except the following disordered-loop regions: M1-G4 (N-terminus), V75-P89, T150-W171, and L262-R313 (C-terminus) (Figure 1C-1E, and Figure S3F-S3H). TWIK-2 assembles as a canonical domain-swapped homodimer with a pseudo-tetrameric central pore observed in all other K_2P_channels (Figure 1). Each TWIK-2 protomer contains four transmembrane helices (M1-M4), two pore-forming helices (PH1 and PH2), two selectivity filter loops (SF1 and SF2), and two extracellular cap-forming helices (CH1 and CH2) (Figure 1C). The cap-forming helices within a protomer are arranged in an inverted ‘V’ shaped cap domain over the central pore of the channel. The cap domains from both protomers form an ‘arched dome,’ creating a bifurcated extracellular ion pathway (EIP). As observed in other known K_2P_ structures, we notice that the cap domain is also responsible for pairing the M1 helix of one protomer in a three-dimensional module with the rest of the protein from the other protomer. Thus, the domain-swapping of the M1 helix leads to the formation of four pore-forming domains comprised of two PD1s that are non-identical to the two PD2s. PD1s comprise the M1 helix from one protomer and PH1, SF1, and M2 from the other protomer. PD2s, on the other hand, are assembled by M3, PH2, SF2, and M4 of the same protomer (Figure 1D-1E).

Three-dimensionally, each pore-forming domain comprises one pore-forming helix (PH) and one selectivity filter (SF) loop flanked by two transmembrane helices. The PH and SF regions from PD1 (PH1/SF1) and PD2 (PH2/SF2) from both protomers form the central ion-conducting pore, wherein M2 and M4 are positioned adjacent to the core, while M1 and M3 are located on the periphery. The domain-swapped M1 helices, the non-identical sequences of the SFs, and unequal lengths of the SF1-M2 and SF2-M4 linkers impart a pseudo-hetero-tetrameric symmetry to the elements surrounding the pore of the channel. Our cryo-EM density map enables the placement of three K^+^ ions in the central selectivity filter (Figure 1D).

### Ion conduction pore of TWIK-2

The ion conduction pathway of TWIK-2 extends from the hydrophobic cuff marked by a kink in the M2 helix within the vestibule designed to funnel ions toward the central pore to the bifurcated EIP on the extracellular side through a narrow selectivity filter (Figure 2A). While the sequence within the selectivity filters of TWIK-2 is largely conserved across different K_2P_ channels, the entry and exit of the ion conductance pathway are flanked by non-conserved M135 and Y111, respectively (Figure S1). We found that the narrowest constriction in the ion conductance pathway has a radius of 4.3 Å, which is wide enough for the K⁺ ions to pass through. We also noticed that the carbonyl oxygens of conserved V107 and Y109 that line the ion permeation pathway face away from the central permeation axis in TWIK-2 (Figure 2B). The backbone oxygens within the SF have been implicated in regulating the K^+^ ion flow. While the SF sequences are primarily conserved across the K_2P_ family, TWIK subfamily members show some divergence. The canonical isoleucine (TIGY/FG) in SF1 is replaced by threonine in TWIK-1 and valine in TWIK-2. Similarly, the canonical phenylalanine (TI/VG<u>F</u>G) in SF2 is replaced by leucine in TWIK-1 and TWIK-2. These variations could govern the differences in the currents of the TWIK subfamily compared to other highly modulated K_2P_ channels, such as TREK and TASK (Figure S1A).

**Figure 2.**
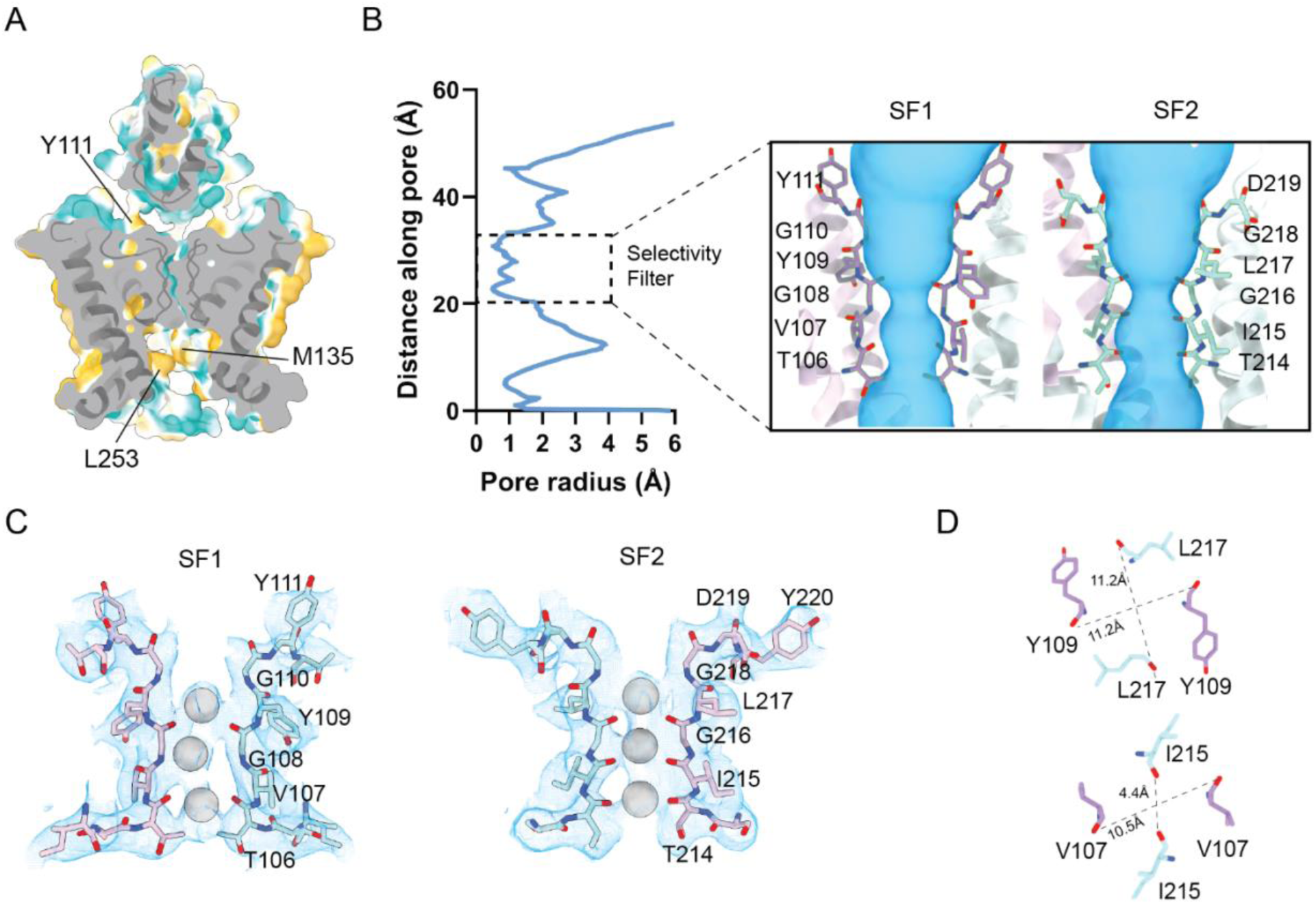
Ion conductance pathway of TWIK-2. **A**. A cross-section of TWIK-2 along the membrane plane, with hydrophobic and hydrophilic amino acids colored in golden yellow and cyan, respectively. Critical amino acids flanking the entry and exit of the ion permeation pore are highlighted. **B**. Pore radius of the TWIK-2 channel as distance along the ion permeation pathway calculated by HOLE (left) ^81^. The ion permeation pore with selectivity filters is highlighted (right). **C.** Cryo-EM density for the selectivity filters and K^+^ ions is highlighted in blue. **D.** The cross-section views from the extracellular cap domain of flipped carbonyls in SF1 and SF2. The distance between the amino acids indicates the pore diameter at that position.

Our automated patch-clamp recordings showed that, unlike TWIK-1, the TWIK-2 channel activity depended on the voltage applied but not the extracellular pH (Figure 1A). Although TWIK-2 and TWIK-1 exhibit high structural similarity, including conserved residues in the SF and M2, subtle differences in residue interactions may lead to different gating behaviors in the two channels. Therefore, we performed site-directed mutagenesis along the ion permeation pathway to identify and assess molecular determinants critical to K⁺ conductance in TWIK-2 (Figure 3A). We mutated two non-conserved residues flanking the entry and exit of the ion conductance pathway (M135 and Y111) and two conserved threonines on the SFs (T106 and T214). While T106 (SF1), T214 (SF2), and M135 (M2) were all mutated to alanine, Y111 (SF1-M2 linker) was mutated to histidine to mimic the corresponding position in TWIK-1 ^39,40^.

**Figure 3.**
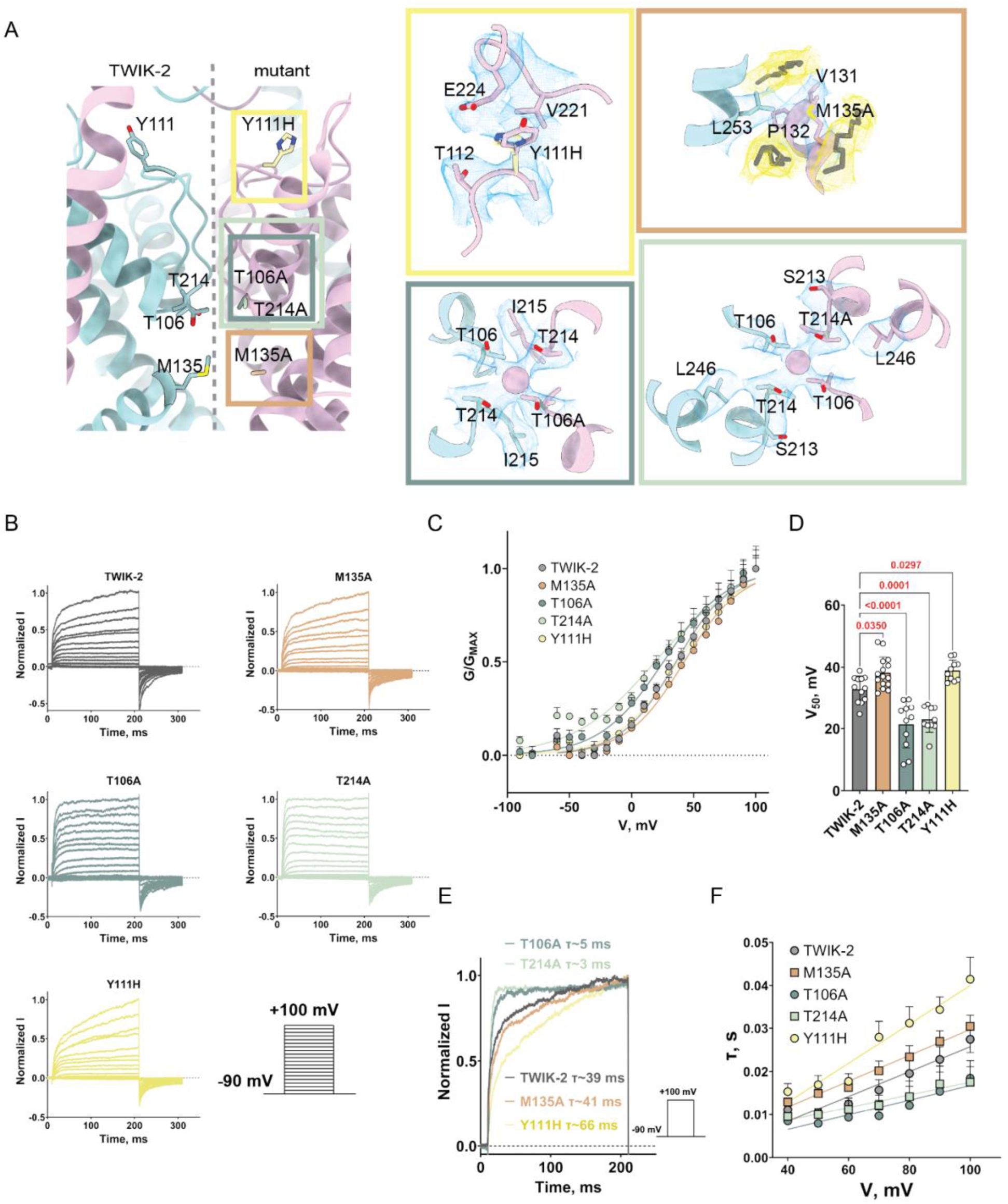
Effects of mutants on TWIK-2 conductance and channel activation. **A**. The relative position of the residues mutated in TWIK-2 along the ion permeation path is highlighted. The insets on the right highlight critical interactions and the amino acid environment of the mutations. The highlights are colored the same way as the mutant data in the remaining figure: T106A (moss), Y111H (yellow), M135A (orange), and T214A (mint). FoldX^82^ was used to generate mutant models. **B.** Channel currents for TWIK-2 and mutants were recorded under high extracellular potassium (K^+^) conditions at pH 7.4. The normalized current traces for both wild-type and mutant TWIK-2 channels are presented. The recording protocol is illustrated in the inset below. **C.** The voltage-dependent activation curves for TWIK-2 (n = 17) and mutant (M135A, n = 18; T106A, n = 12; T214A, n = 10 and Y111H, n = 19) TWIK-2 channels are provided. The data are presented as mean ± SEM. **D.** The half-maximal activation voltage shift (V_50_) for wild-type (n = 12) and mutant (M135A, n = 16; T106A, n = 11; T214A, n = 10 and Y111H, n = 10) TWIK-2 channels is shown, with data expressed as mean ± SD. A one-way ANOVA with Dunnett’s multiple comparisons was performed to evaluate differences between the wild-type and each mutant. **E.** Normalized traces for wild-type and mutant TWIK-2 channels at the activation onset during a 200 ms pulse to +100 mV from a holding potential of −90 mV are displayed. The activation time constants (τ) were determined using a single exponential function**. F**. Activation kinetics were measured for each voltage step in wild-type (n = 28) and mutant channels (M135A, n = 24; T106A, n = 12; T214A, n = 10 and Y111H, n = 37) TWIK-2 channels. The data are expressed as mean ± SEM.

We performed whole-cell recordings of cells expressing wild-type or TWIK-2 channel mutants (Figure 3B). No significant differences in overall current densities were observed in the mutant channels compared to TWIK-2, despite the noticeable differences in the activation profiles of the macroscopic currents (Figure 3B and 3C). However, a closer examination of the activation onset revealed that the Y111H mutation, located at the extracellular mouth of the ion permeation pore, showed delayed activation than TWIK-2 (Figure 3E and 3F). A similar phenomenon was observed for the M135A mutation, located at the entrance of the ion permeation pathway. Still, the increase in activation time was not as pronounced as it was for the Y111H mutation (Figure 3C-3F). In contrast, the mutants T106A and T214A at the selectivity filter (SF1 and SF2, respectively) exhibited faster activation than the wild-type channel (Figure 3E and 3F). We also note a voltage-dependent change in the activation time constants (τ) for the TWIK-2 channel (Figure 3F). For example, the Y111H mutation had a higher activation time constant, implying a slower channel activation. Similarly, the M135A mutant also exhibited slightly slower channel activation. In contrast, T106A and T214A had smaller activation time constants, leading to faster channel activation. Consistent with these findings, Y111H and M135A caused a rightward shift in voltage-dependent activation, G/G_MAX_-V curve, changing the V_50_ from approximately +33 mV to around +39 mV (Figures 3C and 3D). However, T106A and T214A mutants enhanced the voltage sensitivity of TWIK-2 by left-shifting the G/G_MAX_–V curve to approximately +22 mV. We find that alanine mutations of the conserved threonines of the SF (T106A and T214A) that coordinate the canonical site-4 K^+^ ion directly impact the pore domain and enhance gating efficiency, promoting open states over closed ones. In addition, these mutants also alter the time-dependence causing rapid activation of TWIK-2 as evidenced by instantaneous current-voltage responses in the microscopic current responses, compared to the wild-type TWIK-2 or TWIK-2 harboring Y111H and M135A mutations (Figure 3B).

### Modulation of TWIK-2 by small molecules

The unique pseudo-tetrameric architecture of the transmembrane region and the sequence asymmetry between pore-forming domains of the K_2P_ channels are surmised to make them insensitive to existing K^+^ channel blockers. Moreover, the cross-reactivity of known modulators between different sub-families of K_2P_ channels has made efficient targeting of selective K_2P_ channels a bottleneck ^5^. To establish a high-throughput automated approach for screening modulators of TWIK-2, we studied the modulation of TWIK-2 using four known K_2P_ modulators - ML335, BL1249, ML365, and NPBA (Figure 4A). In agreement with previous reports, we observed that ML365 and NPBA inhibit TWIK-2 with IC_50_ values of 1.5 µM and 5.7 µM, respectively (Figure 4B and 4C) ^47,48^. However, we did not see any modulation of TWIK-2 by ML335 and BL1249, which have been shown to modulate TREK and TRAAK family channels (Figure 4B and 4C) ^8,47,53,54^.

**Figure 4.**
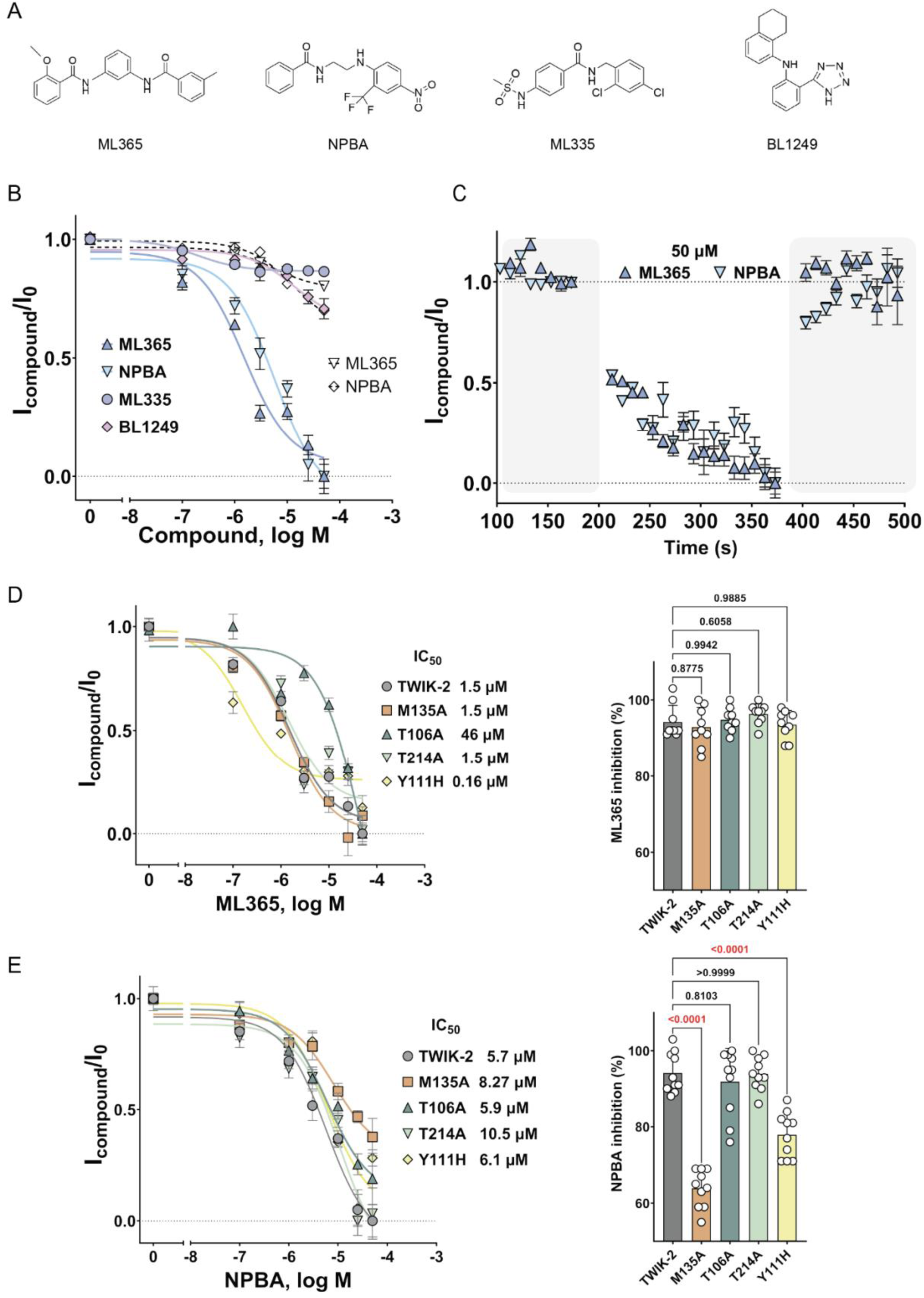
Small molecule modulation of TWIK-2 by known K_2P_ channel inhibitors. **A.** Chemical structure of ML365, NPBA, ML335 and BL1249**. B.** Concentration-response curves of various K2P modulators on TWIK-2 channel inhibition. Data are presented as mean ± SEM: ML365 (n = 8), NPBA (n = 7-8), ML335 (n = 7-8), and BL1249 (n = 7-8). Data from non-T cells (traced lines, empty symbols) are also presented for ML365 (4-7) and NPBA (7-16). **C.** Current responses were measured over time using a voltage-step protocol, applying +60 mV for 350 ms after holding at −90 mV. The results in the blue panel illustrate the changes in TWIK-2 currents in response to ML365 or NPBA (50 μM). Data are presented as the mean ± SEM: ML365 (n = 8), NPBA (n = 8). **D.** Comparison of concentration-response curves of ML365 on wild-type and mutant TWIK-2 channels (left). Data are presented as mean ± SEM: ML365 (n = 8). Scatter dot plots show the current inhibition percentage at 50 µM for ML365 for TWIK-2 (ML365, n = 9) and the mutant channels (right panel, M135A: n = 9; T106A: n = 9; T214A: n = 9; Y111H: n = 9). The data are presented as mean ± SD. A one-way ANOVA with Dunnett’s multiple comparisons test was performed to compare the wild-type channel against each mutant. **E.** Comparison of concentration-response curves of NPBA on wild-type and mutant TWIK-2 channels. Data are presented as mean ± SEM: NPBA (n = 7-8). Right panel is scatter dot plots show the current inhibition percentage at 50 µM NPBA for TWIK-2 (NPBA, n = 10) and the mutant channels (M135A: NPBA, n = 10; T106ANPBA, n = 10; T214A; NPBA, n = 10; Y111H: NPBA, n = 10). The data are presented as mean ± SD. A one-way ANOVA with Dunnett’s multiple comparisons test was performed to compare the wild-type channel against each mutant.

To better understand how ML365 and NPBA inhibit TWIK-2, we compared their effect on the inhibition of TWIK-2 in the context of the ion-permeation pathway mutations (Figure 4D and 4E). In all four mutants, there was negligible effect in the NPBA IC_50_ compared to TWIK-2, albeit there was a reduced maximum inhibition at the highest concentration of NPBA tested in M135A and Y111H. ML365 IC_50_ remained unaltered for M135A and T214A mutants. However, Y111H and T106A variants of TWIK-2 showed enhanced and reduced susceptibility to inhibition by ML365. These findings suggest T106 and Y111 as key determinants of ML365 sensitivity in TWIK-2 and underscore potential differences in the inhibition mechanisms between ML365 and NPBA (Figure 4D and 4E).

## Discussion

Our study employs cryo-electron microscopy (cryo-EM), site-directed mutagenesis, and automated whole-cell patch clamp electrophysiology to reveal insights into the molecular determinants of modulation of TWIK-2 activity. In addition, using known K_2P_ modulators, we established a robust assay to screen for inhibitors specific to TWIK-2.

### Human TWIK-2 in an intermediate conductive state

The ∼3.7 Å cryo-EM structure of TWIK-2 reveals a canonical K_2P_ channel architecture, including the domain-swapped homodimer and a pseudo-tetrameric central pore. However, we also noticed some distinct features in our structure that could be relevant to the mechanism of TWIK-2 action. We observed tubular densities below the SF in the canonical K_2P_ vestibule, modulatory lipid site, and fenestration sites that could not be confidently assigned to the protein (Yellow, Figure 5A). The resolution of our cryo-EM map precludes unambiguous identification of these tubular densities. While we did not supply any exogenous lipids during purification, endogenous lipids are known to co-purify with the protein in the detergent micelle ^55–57^. Alternatively, these densities could be DMNG detergent. However, we do not see densities that can accommodate the head group of DMNG. Instead, we noticed three small tetrahedral blobs within the vestibule below the permeation pore that could accommodate an ion or water (Blue, Figure 5A). Molecular dynamics (MD) simulations of K_2P_ channels treated with negatively charged activators have suggested the presence of ions or water within the “cavity binding sites,” stabilized by the negative moiety of the small molecule or lipids ^58^. We left these small densities unmodeled in our TWIK-2 structure because the precise identity of these densities remains unclear. Based on these observations, we modeled hydrocarbon tails in the tubular densities (Figure 5A).

**Figure 5.**
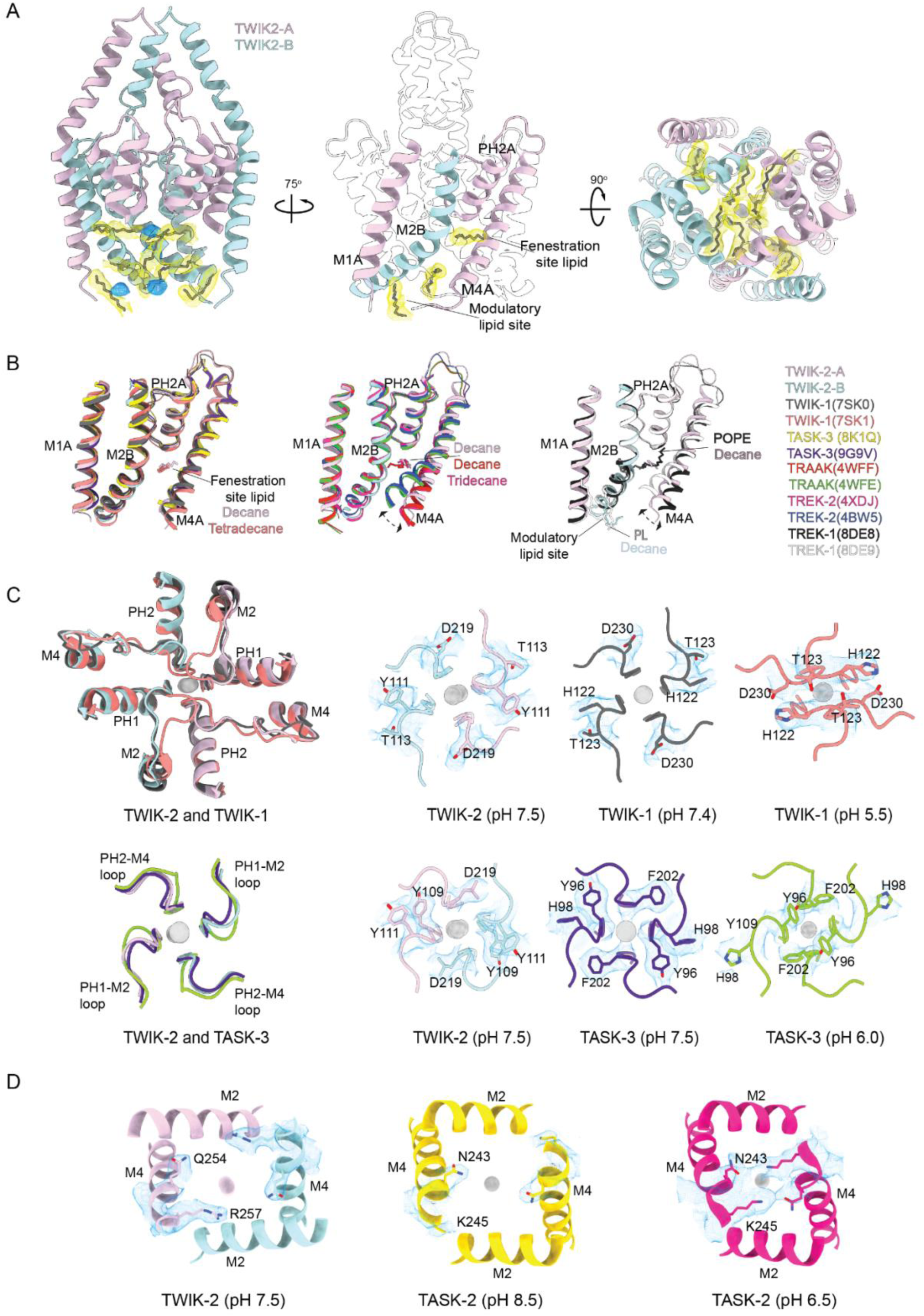
Comparison of TWIK-2 structure with other K2Ps. **A.** TWIK-2 with lipid-like tubular density (yellow) and tetrahedral solvent or ion blobs (blue) highlighted. The center panel highlights the presence of lipid-like densities along the grove made by M1, M4, and PH1 of one protomer with M2 of the other protomer. **B**. A comparison of M4 helix ‘up’ and ‘down’ conformation of different pH-sensitive (first panel) and lipid-regulated K2P channels. (Left) Superposition of the M1A, M2B, PH2A and M4A regions of TWIK-2 (chain A in purple, chain B in cyan) and TWIK-1 at pH 7.4 (black, PDB: 7SK0), TWIK-1 at pH 5.5 (pink, PDB: 7SK1), TASK-3 at pH 6.0 (yellow, PDB: 8K1Q), TASK-3 at pH 7.5 (purple, PDB: 9G9V). Dashed double-arrow lines depict M4 movement. The molecules modeled at the fenestration and modulatory lipid sites have been highlighted in the same color as their respective proteins. (Middle) Superposition of TWIK-2 and non-conductive TRAAK (red, PDB: 4WFF), conductive TRAAK (green, PDB: 4WFE), TREK-2 with M4 in down-state (pink, PDB: 4XDJ) and TREK-2 with M4 in up-state (blue, PDB: 4BW5. (Right) Superposition of TWIK-2, TREK-1 in DDM/POPA mixed micelles (black, PDB: 8DE8), and TREK-1 in DDM/POPE mixed micelles (grey, PDB: 8DE9). **C.** Comparison of the exit of ion permeation pathway of TWIK-2 with pH-sensitive TWIK-1 (top) (pH 7.4 (PDB: 7SK0 and EMDBs: EMD-25168) and pH 5.5 (PDB: 7SK1 and EMD-25159)) and (down) TASK-3 ((pH 7.5 (PDB: 9G9V and EMD-51158) and, pH 6.0 (PDB: 8K1Q and EMD-36799). **D.** Comparison of the entrance of the ion permeation pathway of TWIK-2 (left) with pH-regulated channel TASK-2 in pH 8.5 (yellow, PDB: 6MW0 and EMDB:10422) and in pH 6.5 (bright pink, PDB: 6WLV and EMDB:10423). The cryo-EM densities of critical residues of the hydrogen bond seal from the respective PDBs have been highlighted.

We note that these tubular lipid-like densities occupy the grove created by the PH2 and M4 helix of one protomer with the M2 helix of the other. The lower end of this groove is believed to be a ‘classic’ modulatory phospholipid binding site, and the site that opens to the bottom of SF has been termed a ‘fenestration’ site ^5^. It is postulated that the binding of lipids in this grove pushes the M4 helix down, allowing better access to SF. In a ‘down’ conformation, the M4 helix traverses the lipid bilayer at a ∼45° angle, creating a hydrophobic ‘fenestration site’ between the bottom of the PH2 helix and M2 helix of the opposite protomer that enables binding of hydrophobic compounds at this pocket that inhibit the channel ^5,59^. In TWIK-2, we notice that M4 adopts a canonical ‘down’ conformation explaining the presence of tubular densities in the ‘fenestration site.’

In pH-sensitive channels, such as TWIK-1 and TASK-3, the structures obtained in both open and acid-induced closed conformations show M4 helix in a down conformation, irrespective of bound lipid at the fenestration site ^12,13,35,39^ (Figure 5B). However, in K_2P_ channels that are lipid modulated, such as TREK and TRAAK, only occupation of the fenestration site has been shown to push the M4 helix in a down conformation ^9,15^ (Figure 5B). Lipids at the classic modulatory phospholipid binding site, but not the fenestration site, have been shown to retain the M4 helix in an up conformation ^8,59,60^ (Figure 5B). Perhaps the molecular determinants of pH-induced and lipid-mediated modulation in the K_2P_ channels are different and complex because molecular dynamics simulation studies of TWIK-1 reported that the alkyl tails of surrounding lipids could enter the fenestrations but not far enough to occlude the inner pore ^61^. Notably, despite the ‘down’ conformation of the M4, in our TWIK-2 structure, we observed that the entrance and exit to the ion permeation pore are open, and the SF itself is in an intermediate conductive state.

K^+^ ions exit the ion permeation pore at the space below the cap domain, guarded by SF1-M2 and SF2-M4 loops (Figure 5C). In pH-sensitive channels like TWIK-1 and TASK-3, a conserved histidine residue (H122 in TWIK-1 and H98 in TASK-3) is held towards the transmembrane region inside a hydrophobic pocket between SF1 and PH1. At low pH, protonation of this histidine causes its release from this hydrophobic pocket, rendering the SF1-M2 loop flexible. The histidine either comes together over the ion permeation pore blocking the K^+^ ion exit (TWIK-1) or pulls away, flipping the adjacent bulky aromatic residues into the path of the pore exit (TASK-3) (Figure 5C) ^12,13,35,39^. TWIK-2 has a tyrosine (Y111) compared to the pH-sensing histidine in TWIK-1 and TASK-3. Perhaps this is why TWIK-2 is insensitive to extracellular pH changes in our automated patch-clamp recordings (Figure 1B). Further, we notice that the conformation of SF1-M2 and SF2-M4 loops surrounding the pore in TWIK-2 match open conformations of TWIK-1 and TASK-3. The access to ion permeation pore from the intracellular environment is regulated by a seal formed by residues in the M4 helix (Figure 5D). In TASK-2, another pH-sensitive K_2P_ channel, an asparagine (N243) and lysine (K245), pull the M4 helices over the ion permeation entrance and form a hydrogen bonding network ^27^. K245 side chain is held towards the hydrophobic M2 at basic pH, and at low pH, this residue flips out towards the N243, thereby sealing the entrance to the ion-permeation pathway (Figure 5D). In TWIK-2, we see a similar residue arrangement where R257 is held towards M2 facing away from Q254, precluding the formation of a hydrogen bonding network, thus keeping the entrance to the ion permeation pore open (Figure 5D).

MD simulation studies to understand the role of C-type gating in K_2P_ channels have suggested that extensive conformational dynamics, especially in the SF2-M4 loop and around the fenestration site, are induced by the motion of the M4 helix ^60,62^. It is also believed that movement of the M4 helix from ‘down’ to ‘deep-down’ conformation destabilizes the PH2 helix, causing a perturbation in SF2 and hence causing inactivation of the channel ^62^. In K_v_, Kir K^+^ channels and their prokaryotic homolog (KcsA), C-type inactivation is manifested by perturbations to the carbonyl conformations within the backbone of SF ^63–65^. While the overall conformations of SFs in our structure are largely in a canonical conductive state, we see some minor perturbations (Figure 2B, 2D, and S4). For example, the carbonyl oxygens of Y109 (SF1) and L217 (SF2) around the S0/S1 K^+^ ion site in TWIK-2 face away from the central permeating axis as seen in the case of the equivalent residues in TWIK-1 acid-inhibited low pH structure (Figure S4A and S4B). The positions of carbonyl oxygen of V107 (SF1) around the S2/S3 K^+^ ion site and the side chain of Y111 (SF1) in TWIK-2 don’t match either of the TWIK-1 low and high pH structures, perhaps indicative of sequence variation induced structural differences (Figure S4A and S4B) ^39^.

The canonical TVGYG sequence in the SF1 of TWIK-2 is conserved in homo-tetrameric K⁺-selective channels like KcsA and Kv channels (Figure S1B, Figure S4C). However, most K_2P_ channels deviate from this canonical motif, and this variation has functional and evolutionary implications ^66,67^. Nonetheless, The conformation of SF in KcsA with an E71A mutation in pore-forming helix, which the authors term ‘flipped structure’, is similar to the SF1 in TWIK-2 ^68^. E71A mutant in KcsA has been shown to promote open probability, and the unique SF conformation in the flipped structure has been suggested to be a possible intermediate conductive conformation within the range of possible open states or a transition to the inactivated state. A similar hypothesis is also supported by TWIK-1 simulation studies, which showed that the selectivity filter diverges significantly with conformations that contain defective coordination sites and have been considered partially conductive ^69^. The combination of M4 helix in the ‘down’ conformation, presence of lipid-like tubular densities in the fenestration site, open entrance and exit of the ion permeation pathway, and presence of three K^+^ ions within an SF showing non-canonical carbonyl oxygen positions at S0/S1 and S2/S3 sites, suggest that TWIK-2 structure is in an intermediate conductive state.

### Molecular determinants of K^+^ conductance in TWIK-2

In our automated whole-cell patch-clamp recordings using TWIK-2 expressed in HEK293 GnTI^-^ cells, we notice voltage- and time-dependence of currents, unlike previous observations involving TWIK-1 and TWIK-2 ^70,47^. The currents observed in our experiments are similar to robust TWIK-1 currents recorded using giant inside-out excised oocyte patches, where the currents displayed time- and voltage-dependence of activation as seen in the TREK subfamily of channels ^71^. Perhaps, the lack of native glycosylation of TWIK-2 alters channel properties, including localization, as we used HEK293 GnTI^-^ cells, which lack N-acetylglucosaminyltransferase I activity. Nevertheless, this experimental setup gave us a quick, high-throughput way to measure K^+^ outward currents in TWIK-2 (Figures 1 and 3).

Through site-directed mutagenesis of amino acids along the ion permeation pathway, we identified that conserved amino acids, T106 and T214, act as critical determinants of channel gating in the selectivity filters. The corresponding alanine mutations of these threonines significantly reduced the time- and voltage-dependence of TWIK-2 channel activation (Figure 3B-3F). T106A and T214A are the only conserved amino acids within the SF that coordinate K^+^ ions through their side chains, as opposed to carbonyl oxygens like the remaining amino acids in the SF. T106 in TWIK-2 is also stabilized by interactions with T214 and I215 at the SF2-PH2 interface (Figure 3A). T214 is stabilized by interactions with T106 and S213 around the base of the SF and with L246 of M2 near the hydrophobic cuff. Mutations T106A and T214A disturb these networks of hydrogen bonding and hydrophobic interactions that stabilize the base of the SF and make the channel less selective and more open (Figure 3B-3F). The decrease of time- and voltage-dependence of channel activation in these mutants indicates that recombinant TWIK-2 is voltage-gated. These findings provide a mechanistic framework for understanding TWIK-2 gating. A similar observation was also made for TREK and TASK subfamily channels, where mutating conserved threonine of SF1 and SF2 to cysteine abolished voltage gating ^70^.

The residues at the kink in M2 helix in juxtaposition with M4 helix create a ‘hydrophobic cuff’ that regulates hydration of the central vestibule (Figure 2A and 3A). It is already known that mutation of a single non-polar amino acid at this hydrophobic cuff in K_2P_ channels to a polar or charged residue has been shown to enhance channel activity by promoting hydration of the cavity below the SF ^61,71–73^. We wanted to see if keeping the chemical nature of the sidechain intact by reducing the size of the side chain will influence channel activity. The M135A substitution at the hydrophobic cuff in TWIK-2 does not change the overall channel properties. The M135 sidechain faces L253 of M4 at the hydrophobic cuff and is surrounded by a hydrophobic environment made of lipid-like densities. As a result, replacing the longer non-polar methionine with a shorter non-polar alanine doesn’t drastically alter the channel activity (Figure 3B-3F).

A conserved histidine in the SF1-M2 linker has been shown to be a pH sensor in TWIK-1 and some pH-sensitive TASK family channels. In the structures obtained in basic and acidic pHs, the histidine in the SF1-M2 loop is shown to induce conformational changes near the cap constricting the pore on the extracellular side ^12,13,35,39,40^. While the residues surrounding this histidine are conserved in TWIK-2, histidine itself is replaced by a tyrosine (Figure 3A). Hence, we see that TWIK-2 is not pH sensitive. While our Y111H mutation shows increase in the time taken for activation, perhaps because of the interaction of the introduced histidine at this position with E224 of the SF2-M4 linker preventing the required conformational freedom of the SF1-M2 and SF2-M4 loops around the ion permeation exit (Figure 3).

### Small molecule modulation of TWIK-2

High-throughput screening of K_2P_ channel modulators has been previously reported, especially by assaying the growth of engineered yeast whose survival depended on the expression of functional K_2P_ channels ^22,74^. Here, we confirm the feasibility of using an automated whole-cell patch-clamp electrophysiology approach as a high-throughput platform to screen for TWIK-2 modulators in HEK cells. We used four known K_2P_ modulators in our trial – ML335, BL1249, NPBA, and ML365 – and noticed that we could see inhibition by only NPBA and ML365 (Figure 4A and 4B). In the case of NPBA, we do not see a prominent change in the IC_50_ of inhibition, except for a mild decrease in the efficacy in T214A and M135A, both mutations that face fenestration site and central vestibule (Figure 3 and 4E). Conversely, in the case of ML365, we do not see any difference in the IC_50_ for T214A and M135A, but we see an altered dose response for T106A and Y111H. Interestingly, the change in IC_50_ for T106A and Y111H are opposite. Y111H improves the efficacy of ML365 while T106A reduces it, although both mutations reside on SF1, albeit on either end of it. As noted earlier, Y111H potentially minimizes the movement of the SF2-M4 and SF1-M2 linkers by interacting with E224. Combined with a reduced steric clash because of Y-H substitution, this reduced flexibility could allow for better binding of ML365. We see a decreased efficacy for T106A but not for T214A, perhaps because PD1 asymmetrically governs the binding of ML365 as opposed to PD2, as both Y111H and T106A are a part of PD1. Conversely, the differential effect of ML365 in mutations that reside on either end of SF1 could indicate an allosteric coupling between the two sides of SF. However, we do not have clear evidence to propose a binding site for ML365 on TWIK-2 because we also note that in ΔC-TWIK-2, we see a significant reduction in percentage inhibition at maximum drug concentration, in the case of ML365 but not NPBA (Figure S2E). Nonetheless, our studies suggest a potential difference in TWIK-2 binding sites and mechanism of action for ML365 and NPBA.

## Conclusion

The intermediate conductive state of TWIK-2 raises intriguing questions about its regulation. Given the pH-independent and voltage-dependent findings in our electrophysiology experiments and the presence of lipid-like densities at the modulatory lipid binding site and fenestration site in our cryo-EM map, identifying cellular signals that modulate the activity of TWIK-2 becomes pertinent. Furthermore, the weak (micromolar) IC_50_ of the known K_2P_ channel inhibitors underscores the need for potent and selective modulators of TWIK-2. In this study, we characterize the structural and functional properties of TWIK-2 and offer essential insights into its ion selectivity, gating mechanisms, and modulation by small molecules of TWIK-2. These results enhance our understanding of TWIK-2 within the larger context of K_2P_ channels and lay a solid foundation for future research to target TWIK-2. Our structure and the automated whole-cell patch-clamp platform provide a robust start for structure-based design and high-throughput screening of modulators with improved efficacy and specificity.

## Data availability

The density map of TWIK-2 has been deposited in the Electron Microscopy Data Bank under the accession number EMD-47768. 3D coordinates for the TWIK-2 structure have been deposited in the Protein Data Bank (PDB) under the accession code 9E94.

## Author contributions

VN and SM designed the study. QM, AL, and VN prepared the sample. QM collected the cryo-EM data. QM, VN, and AK processed the cryo-EM data. AK and VN built the model and deposited the coordinates. CCH performed all electrophysiology experiments. QM, CCH, VN, and SM analyzed the data. VN, QM, and CCH wrote the manuscript. All authors contributed to finalizing the manuscript.

## Acknowledgments

We appreciate the efforts of the staff at the University of Michigan (U-M) cryo-EM facility in assisting with cryo-EM data collection. We thank Dr. Sandipan Chowdhury at the Department of Molecular Physiology and Biophysics, University of Iowa, for helpful discussions on our work. We also thank members of the Mosalaganti lab and the Baldridge lab for their comments on our work. Jiameng Zong and Jaimin Rana are acknowledged for their assistance in cell culturing and sample preparation. We thank U-M BSI and U-M LSI for their generous support of the U-M cryo-EM facility. The Nanion Synchropatch used for patch clamp recording was supported by an NIH grant (1S10OD025203) to the Center for Chemical Genomics at the U-M. The Klatskin-Sutker Discovery Fund award funded a portion of the research presented here to QM, and the start-up funding was provided to SM by the U-M Life Sciences Institute.

## References

1. Lesage, F., and Lazdunski, M. (2000). Molecular and functional properties of two-pore-domain potassium channels. Am. J. Physiol. Renal Physiol. 279, F793–F801.

2. O’Connell, A.D., Morton, M.J., and Hunter, M. (2002). Two-pore domain K+ channels-molecular sensors. Biochim. Biophys. Acta 1566, 152–161.

3. Buckingham, S.D., Kidd, J.F., Law, R.J., Franks, C.J., and Sattelle, D.B. (2005). Structure and function of two-pore-domain K+ channels: contributions from genetic model organisms. Trends Pharmacol. Sci. 26, 361–367.

4. Enyedi, P., and Czirják, G. (2010). Molecular background of leak K+ currents: two-pore domain potassium channels. Physiol. Rev. 90, 559–605.

5. Natale, A.M., Deal, P.E., and Minor, D.L., Jr (2021). Structural insights into the mechanisms and pharmacology of K2P potassium channels. J. Mol. Biol. 433, 166995.

6. Feliciangeli, S., Chatelain, F.C., Bichet, D., and Lesage, F. (2015). The family of K2P channels: salient structural and functional properties: The family of K_2P_channels. J. Physiol. 593, 2587–2603.

7. Mathie, A., and Veale, E.L. (2015). Two-pore domain potassium channels: potential therapeutic targets for the treatment of pain. Pflugers Arch. 467, 931–943.

8. Lolicato, M., Arrigoni, C., Mori, T., Sekioka, Y., Bryant, C., Clark, K.A., and Minor, D.L., Jr (2017). K2P2.1 (TREK-1)-activator complexes reveal a cryptic selectivity filter binding site. Nature 547, 364–368.

9. Dong, Y.Y., Pike, A.C.W., Mackenzie, A., McClenaghan, C., Aryal, P., Dong, L., Quigley, A., Grieben, M., Goubin, S., Mukhopadhyay, S., et al. (2015). K2P channel gating mechanisms revealed by structures of TREK-2 and a complex with Prozac. Science 347, 1256–1259.

10. Brohawn, S.G., del Mármol, J., and MacKinnon, R. (2012). Crystal structure of the human K2P TRAAK, a lipid- and mechano-sensitive K+ ion channel. Science 335, 436–441.

11. Rödström, K.E.J., Kiper, A.K., Zhang, W., Rinné, S., Pike, A.C.W., Goldstein, M., Conrad, L.J., Delbeck, M., Hahn, M.G., Meier, H., et al. (2020). A lower X-gate in TASK channels traps inhibitors within the vestibule. Nature 582, 443–447.

12. Lin, H., Li, J., Zhang, Q., Yang, H., and Chen, S. (2024). C-type inactivation and proton modulation mechanisms of the TASK3 channel. Proc. Natl. Acad. Sci. U. S. A. 121, e2320345121.

13. Hall, P.R., Jouen-Tachoire, T., Schewe, M., Proks, P., Baukrowitz, T., Carpenter, E.P., Newstead, S., Rödström, K.E.J., and Tucker, S.J. (2025). Structures of TASK-1 and TASK-3 K2P channels provide insight into their gating and dysfunction in disease. Structure 33, 115–122.e4.

14. Brohawn, S.G. (2015). How ion channels sense mechanical force: insights from mechanosensitive K2P channels TRAAK, TREK1, and TREK2. Ann N Y Acad Sci 1352, 20–32.

15. Brohawn, S.G., Su, Z., and MacKinnon, R. (2014). Mechanosensitivity is mediated directly by the lipid membrane in TRAAK and TREK1 K+ channels. Proc. Natl. Acad. Sci. U. S. A. 111, 3614–3619.

16. Brohawn, S.G., Campbell, E.B., and MacKinnon, R. (2014). Physical mechanism for gating and mechanosensitivity of the human TRAAK K+ channel. Nature 516, 126–130.

17. Rietmeijer, R.A., Sorum, B., Li, B., and Brohawn, S.G. (2021). Physical basis for distinct basal and mechanically gated activity of the human K+ channel TRAAK. Neuron 109, 2902–2913.e4.

18. Sorum, B., Docter, T., Panico, V., Rietmeijer, R., and Brohawn, S. (2024). Tension activation of mechanosensitive two-pore domain K+ channels TRAAK, TREK-1, and TREK-2. Nat. Commun. 15, 3142.

19. Vivier, D., Soussia, I.B., Rodrigues, N., Lolignier, S., Devilliers, M., Chatelain, F.C., Prival, L., Chapuy, E., Bourdier, G., Bennis, K., et al. (2017). Development of the first two-pore domain potassium channel TWIK-related K+ channel 1-selective agonist possessing in vivo antinociceptive activity. J. Med. Chem. 60, 1076–1088.

20. Pope, L., Lolicato, M., and Minor, D.L., Jr (2020). Polynuclear ruthenium amines inhibit K2P channels via a “finger in the dam” mechanism. Cell Chem. Biol. 27, 511–524.e4.

21. Bagriantsev, S.N., Peyronnet, R., Clark, K.A., Honoré, E., and Minor, D.L., Jr (2011). Multiple modalities converge on a common gate to control K2P channel function: A common gate controls K2P function. EMBO J. 30, 3594–3606.

22. Bagriantsev, S.N., Ang, K.-H., Gallardo-Godoy, A., Clark, K.A., Arkin, M.R., Renslo, A.R., and Minor, D.L., Jr (2013). A high-throughput functional screen identifies small molecule regulators of temperature- and mechano-sensitive K2P channels. ACS Chem. Biol. 8, 1841–1851.

23. Dadi, P.K., Vierra, N.C., Days, E., Dickerson, M.T., Vinson, P.N., Weaver, C.D., and Jacobson, D.A. (2017). Selective small molecule activators of TREK-2 channels stimulate dorsal root ganglion c-fiber nociceptor two-pore-domain potassium channel currents and limit calcium influx. ACS Chem. Neurosci. 8, 558–568.

24. Tian, F., Qiu, Y., Lan, X., Li, M., Yang, H., and Gao, Z. (2019). A small-molecule compound selectively activates K2P channel TASK-3 by acting at two distant clusters of residues. Mol. Pharmacol. 96, 26–35.

25. Alloui, A., Zimmermann, K., Mamet, J., Duprat, F., Noël, J., Chemin, J., Guy, N., Blondeau, N., Voilley, N., Rubat-Coudert, C., et al. (2006). TREK-1, a K+ channel involved in polymodal pain perception. EMBO J. 25, 2368–2376.

26. Maingret, F., Fosset, M., Lesage, F., Lazdunski, M., and Honoré, E. (1999). TRAAK is a mammalian neuronal mechano-gated K+ channel. J. Biol. Chem. 274, 1381–1387.

27. Li, B., Rietmeijer, R.A., and Brohawn, S.G. (2020). Structural basis for pH gating of the two-pore domain K+ channel TASK2. Nature 586, 457–462.

28. Li, X.-Y., and Toyoda, H. (2015). Role of leak potassium channels in pain signaling. Brain Res. Bull. 119, 73–79.

29. Mathie, A., Rees, K.A., El Hachmane, M.F., and Veale, E.L. (2010). Trafficking of neuronal two pore domain potassium channels. Curr. Neuropharmacol. 8, 276–286.

30. Lesage, F., Guillemare, E., Fink, M., Duprat, F., Lazdunski, M., Romey, G., and Barhanin, J. (1996). TWIK-1, a ubiquitous human weakly inward rectifying K+ channel with a novel structure. EMBO J. 15, 1004–1011.

31. Chavez, R.A., Gray, A.T., Zhao, B.B., Kindler, C.H., Mazurek, M.J., Mehta, Y., Forsayeth, J.R., and Yost, C.S. (1999). TWIK-2, a new weak inward rectifying member of the tandem pore domain potassium channel family. J. Biol. Chem. 274, 7887–7892.

32. Pountney, D.J., Gulkarov, I., Vega-Saenz de Miera, E., Holmes, D., Saganich, M., Rudy, B., Artman, M., and Coetzee, W.A. (1999). Identification and cloning of TWIK-originated similarity sequence (TOSS): a novel human 2-pore K+ channel principal subunit. FEBS Lett. 450, 191–196.

33. Lloyd, E.E., Marrelli, S.P., Namiranian, K., and Bryan, R.M., Jr (2009). Characterization of TWIK-2, a two-pore domain K+ channel, cloned from the rat middle cerebral artery. Exp. Biol. Med. (Maywood) 234, 1493–1502.

34. Bobak, N., Feliciangeli, S., Chen, C.-C., Ben Soussia, I., Bittner, S., Pagnotta, S., Ruck, T., Biel, M., Wahl-Schott, C., Grimm, C., et al. (2017). Recombinant tandem of pore-domains in a Weakly Inward rectifying K+ channel 2 (TWIK2) forms active lysosomal channels. Sci. Rep. 7, 649.

35. Miller, A.N., and Long, S.B. (2012). Crystal structure of the human two-pore domain potassium channel K2P1. Science 335, 432–436.

36. Rajan, S., Plant, L.D., Rabin, M.L., Butler, M.H., and Goldstein, S.A.N. (2005). Sumoylation silences the plasma membrane leak K+ channel K2P1. Cell 121, 37–47.

37. Feliciangeli, S., Bendahhou, S., Sandoz, G., Gounon, P., Reichold, M., Warth, R., Lazdunski, M., Barhanin, J., and Lesage, F. (2007). Does sumoylation control K2P1/TWIK1 background K+ channels? Cell 130, 563–569.

38. Chatelain, F.C., Bichet, D., Douguet, D., Feliciangeli, S., Bendahhou, S., Reichold, M., Warth, R., Barhanin, J., and Lesage, F. (2012). TWIK1, a unique background channel with variable ion selectivity. Proc. Natl. Acad. Sci. U. S. A. 109, 5499–5504.

39. Turney, T.S., Li, V., and Brohawn, S.G. (2022). Structural Basis for pH-gating of the K+ channel TWIK1 at the selectivity filter. Nat. Commun. 13, 3232.

40. Chatelain, F.C., Gilbert, N., Bichet, D., Jauch, A., Feliciangeli, S., Lesage, F., and Bignucolo, O. (2024). Mechanistic basis of the dynamic response of TWIK1 ionic selectivity to pH. Nat. Commun. 15, 3849.

41. Patel, A.J., Maingret, F., Magnone, V., Fosset, M., Lazdunski, M., and Honoré, E. (2000). TWIK-2, an inactivating 2P domain K+ channel. J. Biol. Chem. 275, 28722–28730.

42. Mhatre, A.N., Li, J., Chen, A.F., Yost, C.S., Smith, R.J.H., Kindler, C.H., and Lalwani, A.K. (2004). Genomic structure, cochlear expression, and mutation screening of KCNK6, a candidate gene for DFNA4. J. Neurosci. Res. 75, 25–31.

43. Pandit, L.M., Lloyd, E.E., Reynolds, J.O., Lawrence, W.S., Reynolds, C., Wehrens, X.H.T., and Bryan, R.M. (2014). TWIK-2 channel deficiency leads to pulmonary hypertension through a rho-kinase-mediated process. Hypertension 64, 1260–1265.

44. Yu, J., Fu, Y., Zhang, N., Gao, J., Zhang, Z., Jiang, X., Chen, C., and Wen, Z. (2024). Extracellular histones promote TWIK2-dependent potassium efflux and associated NLRP3 activation in alveolar macrophages during sepsis-induced lung injury. Inflamm. Res. 73, 1137–1155.

45. Di, A., Xiong, S., Ye, Z., Malireddi, R.K.S., Kometani, S., Zhong, M., Mittal, M., Hong, Z., Kanneganti, T.-D., Rehman, J., et al. (2018). The TWIK2 potassium efflux channel in macrophages mediates NLRP3 inflammasome-induced inflammation. Immunity 49, 56– 65.e4.

46. Huang, L.S., Anas, M., Xu, J., Zhou, B., Toth, P.T., Krishnan, Y., Di, A., and Malik, A.B. (2023). Endosomal trafficking of two-pore K+ efflux channel TWIK2 to plasmalemma mediates NLRP3 inflammasome activation and inflammatory injury. Elife 12, e83842.

47. Wu, X.-Y., Lv, J.-Y., Zhang, S.-Q., Yi, X., Xu, Z.-W., Zhi, Y.-X., Zhao, B.-X., Pang, J.-X., Yung, K.K.L., Liu, S.-W., et al. (2022). ML365 inhibits TWIK2 channel to block ATP-induced NLRP3 inflammasome. Acta Pharmacol. Sin. 43, 992–1000.

48. Zhi, Y., Wu, X., Chen, Y., Chen, X., Chen, X., Luo, H., Yi, X., Lin, X., Ma, L., Chen, Y., et al. (2023). A novel TWIK2 channel inhibitor binds at the bottom of the selectivity filter and protects against LPS-induced experimental endotoxemia in vivo. Biochem. Pharmacol. 218, 115894.

49. Djillani, A., Mazella, J., Heurteaux, C., and Borsotto, M. (2019). Role of TREK-1 in health and disease, focus on the central nervous system. Front. Pharmacol. 10, 379.

50. Ávalos Prado, P., Chassot, A.-A., Landra-Willm, A., and Sandoz, G. (2022). Regulation of two-pore-domain potassium TREK channels and their involvement in pain perception and migraine. Neurosci. Lett. 773, 136494.

51. Landra-Willm, A., Karapurkar, A., Duveau, A., Chassot, A.A., Esnault, L., Callejo, G., Bied, M., Häfner, S., Lesage, F., Wdziekonski, B., et al. (2023). A photoswitchable inhibitor of TREK channels controls pain in wild-type intact freely moving animals. Nat. Commun. 14, 1160.

52. Mendez-Otalvaro, E., Kopec, W., and de Groot, B.L. (2024). Effect of two activators on the gating of a K2P channel. Biophys. J. 0. 123, 3408–3420

53. Song, D., Zhou, X., Yu, Q., Li, R., Dai, Q., and Zeng, M. (2024). ML335 inhibits TWIK2 channel-mediated potassium efflux and attenuates mitochondrial damage in MSU crystal-induced inflammation. J. Transl. Med. 22, 785.

54. Pope, L., Arrigoni, C., Lou, H., Bryant, C., Gallardo-Godoy, A., Renslo, A.R., and Minor, D.L., Jr (2018). Protein and chemical determinants of BL-1249 action and selectivity for K2P channels. ACS Chem. Neurosci. 9, 3153–3165.

55. Jandu, R.S., Yu, H., Zhao, Z., Le, H.T., Kim, S., Huan, T., and Duong van Hoa, F. (2024). Capture of endogenous lipids in peptidiscs and effect on protein stability and activity. iScience 27, 109382.

56. Dörr, J.M., Koorengevel, M.C., Schäfer, M., Prokofyev, A.V., Scheidelaar, S., van der Cruijsen, E.A.W., Dafforn, T.R., Baldus, M., and Killian, J.A. (2014). Detergent-free isolation, characterization, and functional reconstitution of a tetrameric K+ channel: the power of native nanodiscs. Proc. Natl. Acad. Sci. U. S. A. 111, 18607–18612.

57. Palsdottir, H., and Hunte, C. (2004). Lipids in membrane protein structures. Biochim. Biophys. Acta 1666, 2–18.

58. Schewe, M., Sun, H., Mert, Ü., Mackenzie, A., Pike, A.C.W., Schulz, F., Constantin, C., Vowinkel, K.S., Conrad, L.J., Kiper, A.K., et al. (2019). A pharmacological master key mechanism that unlocks the selectivity filter gate in K+ channels. Science 363, 875–880.

59. Schmidpeter, P.A.M., Petroff, J.T., 2nd, Khajoueinejad, L., Wague, A., Frankfater, C., Cheng, W.W.L., Nimigean, C.M., and Riegelhaupt, P.M. (2023). Membrane phospholipids control gating of the mechanosensitive potassium leak channel TREK1. Nat. Commun. 14, 1077.

60. Lolicato, M., Natale, A.M., Abderemane-Ali, F., Crottès, D., Capponi, S., Duman, R., Wagner, A., Rosenberg, J.M., Grabe, M., and Minor, D.L., Jr (2020). K2P channel C-type gating involves asymmetric selectivity filter order-disorder transitions. Sci. Adv. 6, eabc9174.

61. Aryal, P., Abd-Wahab, F., Bucci, G., Sansom, M.S.P., and Tucker, S.J. (2015). Influence of lipids on the hydrophobic barrier within the pore of the TWIK-1 K2P channel. Channels (Austin) 9, 44–49.

62. Zhang, Q., Fu, J., Zhang, S., Guo, P., Liu, S., Shen, J., Guo, J., and Yang, H. (2022). “C-type” closed state and gating mechanisms of K2P channels revealed by conformational changes of the TREK-1 channel. J. Mol. Cell Biol.14(1), mjac002

63. Yellen, G., Sodickson, D., Chen, T.Y., and Jurman, M.E. (1994). An engineered cysteine in the external mouth of a K+ channel allows inactivation to be modulated by metal binding. Biophys J 66, 1068–1075.

64. Schulte, U., Weidemann, S., Ludwig, J., Ruppersberg, J., and Fakler, B. (2001). K(+)- dependent gating of K(ir)1.1 channels is linked to pH gating through a conformational change in the pore. J Physiol 534, 49–58.

65. Cuello, L.G., Jogini, V., Cortes, D.M., and Perozo, E. (2010). Structural mechanism of C-type inactivation in K(+) channels. Nature 466, 203–208.

66. Andrini, O., Ben Soussia, I., Tardy, P., Walker, D.S., Peña-Varas, C., Ramírez, D., Gendrel, M., Mercier, M., El Mouridi, S., Leclercq-Blondel, A., et al. (2024). Constitutive sodium permeability in a C. elegans two-pore domain potassium channel. Proc. Natl. Acad. Sci. U. S. A. 121, e2400650121.

67. González, W., Valdebenito, B., Caballero, J., Riadi, G., Riedelsberger, J., Martínez, G., Ramírez, D., Zúñiga, L., Sepúlveda, F.V., Dreyer, I., et al. (2015). K₂p channels in plants and animals. Pflugers Arch. 467, 1091–1104.

68. Cordero-Morales, J.F., Cuello, L.G., Zhao, Y., Jogini, V., Cortes, D.M., Roux, B., and Perozo, E. (2006). Molecular determinants of gating at the potassium-channel selectivity filter. Nat. Struct. Mol. Biol. 13, 311–318.

69. Oakes, V., Furini, S., Pryde, D., and Domene, C. (2016). Exploring the dynamics of the TWIK-1 channel. Biophys. J. 111, 775–784.

70. Schewe, M., Nematian-Ardestani, E., Sun, H., Musinszki, M., Cordeiro, S., Bucci, G., de Groot, B.L., Tucker, S.J., Rapedius, M., and Baukrowitz, T. (2016). A non-canonical voltage-sensing mechanism controls gating in K2P K(+) channels. Cell 164, 937–949.

71. Nematian-Ardestani, E., Abd-Wahab, F., Chatelain, F.C., Sun, H., Schewe, M., Baukrowitz, T., and Tucker, S.J. (2020). Selectivity filter instability dominates the low intrinsic activity of the TWIK-1 K2P K+ channel. J. Biol. Chem. 295, 610–618.

72. Ben Soussia, I., El Mouridi, S., Kang, D., Leclercq-Blondel, A., Khoubza, L., Tardy, P., Zariohi, N., Gendrel, M., Lesage, F., Kim, E.-J., et al. (2019). Mutation of a single residue promotes gating of vertebrate and invertebrate two-pore domain potassium channels. Nat. Commun. 10, 787.

73. Poudel, B., Tatikonda, R.R., and Vanegas, J.M. (2019). Exploring the hydrophobic barrier of human K2P channel TWIK1 with steered MD simulations. Biophys. J. 116, 208a.

74. Su, Z., Brown, E.C., Wang, W., and MacKinnon, R. (2016). Novel cell-free high-throughput screening method for pharmacological tools targeting K+ channels. Proc. Natl. Acad. Sci. U. S. A. 113, 5748–5753.

