## Supplementary Material for "Insights into the structure and modulation of human TWIK-2"

**The PDF file includes:**

Materials and Methods

Figs. S1-S4

Table S1

References (75-85)

#### Materials and Methods

##### Cloning and overexpression of recombinant TWIK-2

The codon-optimized full-length wild-type human TWIK-2 (*KCNK6*) was synthesized by GenScript and was subcloned into the pEG BacMam expression vector (Addgene plasmid #160683) to be expressed with an N-terminal Strep-tag-II, followed by an HRV-3C protease site and a GFP tag. TWIK-2 mutants (T106A, Y111H, M135A, T214A) and a deletion construct lacking amino acids 277-313 on the C-terminus (ΔC-TWIK-2) were also made in this background by GenScript. Wild-type (TWIK-2) and variants of TWIK-2 were expressed in HEK293 GnTI- cells (ATCC # CRL-3022). Large-scale protein production was done using a baculoviral expression system as per the protocol described by Goehring et al. [75]. Briefly, the TWIK-2 expression vector was transformed into chemically competent DH10Bac cells (Thermo Fisher Scientific), and white colonies were selected on gentamicin-kanamycin-tetracycline LB agar plates. Isolated Bacmid DNA was used to transfect sf9 cells (ATCC #12659017) with Cellfectin-II reagent (Gibco) per the manufacturer's protocol to make baculovirus P1. P1 was then amplified to make P2 by adding 200 µl of P1 to 200 ml culture of sf9 cells at a $1x10^6$ cells/ml density. Cells were grown at 27℃ for 96 h and harvested by spinning at 4000 x*g* for 20 min. The supernatant was filtered with a 0.22 µm filter and stored at 4℃ in the dark as a P2 virus. This was subsequently used for large-scale transduction of HEK293 $GnTI^-$ suspension adapted cells grown in FreeStyle 293 media supplemented with 2% heat-inactivated fetal bovine serum (FBS, Gibco) at density 2.5-3.0 $x10^6$ cells/ml. Transduced cells were incubated on an orbital shaker at 37℃ and 8% $CO_2$ at 120 rpm. Eight hours after transduction, 10 mM Sodium butyrate was added, and cells were grown for an additional 16-24 h at 37℃, 8% $CO_2$. Cells were harvested by centrifugation at 4000 x*g* for 20 min, flash-frozen in liquid nitrogen, and stored at -80℃.

##### Purification of recombinant human TWIK-2

Cell pellets from 2.4L culture were thawed and lysed in 120 ml lysis buffer (25 mM Tris-HCl, 150 mM KCl, 0.8 µM aprotinin, 2 µg/ml leupeptin, 2 µM pepstatin A, 1 mM PMSF, pH 7.5) by sonication. Lysed cells were centrifuged at 2400 xg for 5 min to remove cell debris. The supernatant was then ultracentrifuged at 185,000 xg for 45 min. The pellet from ultracentrifugation was resuspended using a Dounce homogenizer in 60 ml of lysis buffer. To solubilize the membrane, 60 ml 2X solubilization buffer (25 mM Tris-HCl, 150 mM KCl, 0.8 µM aprotinin, 2 ug/ml leupeptin, 2 µM pepstatin A, 1 mM PMSF, 2% DMNG, pH 7.5) was added and the solubilization was carried out for 90 min in 4℃. The solubilized membrane was then centrifuged at 185,000 xg for 1h. The supernatant was filtered through a 0.22 um filter and was loaded onto a column containing 3 ml Strep-Tactin resin (IBA Life Sciences) at a flow rate of 0.5 ml/min. The column was then washed with 15 ml of wash buffer (25 mM Tris-HCl, 150 mM KCl, 0.1% DMNG, pH 7.5) and eluted with 12 ml elution buffer (25 mM Tris-HCl, 150 mM KCl, 0.1% DMNG, 5 mM Desthiobiotin, pH 7.5) in 1.5 ml fractions. The homogeneity of eluted fractions was analyzed by SDS-PAGE and subsequently pooled and concentrated to 7 mg/ml. In-house purified HRV 3C protease was added to purified TWIK-2 at a molar ratio of 1:20 and incubated overnight at 4℃ to remove the GFP tag. The digested protein was further purified by size-exclusion chromatography using Superdex 200 Increase 10/300 GL column (Cytiva) pre-equilibrated with SEC buffer (0.1%

DMNG, 25 mM Tris-HCl, 150 mM KCl, pH 7.5). The fractions corresponding to GFP-tag-free TWIK-2 dimer were pooled and concentrated to 2.3 mg/ml for cryo-EM sample preparation.

##### Cryo-EM sample preparation and data collection

UltraAuFoil 300 mesh 1.2/1.3 grids (Quantifoil) were glow discharged (PELCO) for 60 s at 15 mA before use. 3.5 µl purified TWIK-2 was applied to the glow-discharged grids and blotted for 2.5 - 3s at $4^{o}$C and 100% humidity using a Vitrobot (Mark IV, ThermoFisher Scientific). Grids were immediately plunged-frozen in liquid ethane. Grids were screened, and movies were collected at the University of Michigan cryo-EM facility on 300 kV Titan Krios G4i cryo-EM (Thermo Fisher Scientific) equipped with K3 direct detector camera (Gatan), and a post-BioQuantum GIF energy filter (Gatan), slit width set to 20 eV. 15,006 movies were collected using SerialEM 4.0 software in counting mode at a magnification of 105,000x (nominal pixel size of 0.83 Å/pixel). The total dose was 60 $e^{-}$ $Å^{-2}$ with 60 frames with a defocus range between 0.8~2.5 µm.

##### Cryo-EM data processing

Data processing was performed in cryoSPARC (version 4.5.3)[76] Briefly, 15,006 movie stacks were motion corrected and patch-CTF estimated, followed by a manual curation to remove micrographs with relatively thick ice and CTF fit > 7 Å. 10,629 micrographs were subsequently selected and used for reference-free particle picking by blob picker with minimum particle diameter of 80 Å and maximum particle diameter of 100 Å followed by extraction at a box size of 256 pixels at binning of 2. 5,447,554 particles were picked and subjected to two rounds of 2D classification that resulted in 1,387,475 relative particles. A subset of ~380,000 particles from 2D classes with clear secondary structure features were used to generate four *ab-initio* 3D classes with C1 symmetry. One "good" class with clear transmembrane features and one "bad" class representing empty micelle were selected as reference input for downstream heterogenous refinement with the total relatively clean 1,387,475 particles. Multiple rounds of heterogeneous refinement were performed until no further improvement of the resolution. 301,000 selected particles from the best class after the last heterogenous refinement were re-extracted with 256 pixels and subjected to one round of Non-uniform refinement with C1 symmetry. A 3D mask was generated using volume of the best *ab-initio class* in Chimera 1.17.1 by manually removing the micelle density and imported into cryoSPARC with a threshold of 0.064 and dilation radius of 3 pix. This mask was used for a focused 3D classification of the clean re-extracted particles into 4 classes in cryoSPARC. 104,997 particles from the class with clear secondary structure features were selected for reference-based motion correction, and refined by non-uniform refinement to obtain a final map at ~3.7 Å.

##### Model building, refinement, and structure analysis

The sharpened map from the final non-uniform refinement in cryoSPARC was used to build a preliminary model in ModelAngelo[77]. The preliminary model was analyzed and improperly built regions were rebuilt manually in Coot [78]. Final structure refinement was performed in Phenix [79]. Figures were prepared using UCSF ChimeraX [80]. The radii along the channel pore was calculated using HOLE [81]. FoldX was used to generate mutant models of TWIK-2, and LigPlot+ was used to review the mutant environment analysis [82,83].

**Automated patch clamp recordings**

The wild-type and mutant TWIK-2 expression constructs were used to transfect adherent HEK293 GnTI$^-$ cells for automated whole-cell patch clamp recordings. Briefly, 10 x $10^6$ cells grown in DMEM media with 10% FBS were transfected, at approximately 1 x $10^6$ cells/ml density, with 4 µg of DNA using TurboFect (Thermo Fisher Scientific) suspended in serum-free DMEM as suggested by the manufacturer. After 8h, the media was replaced with fresh DMEM containing 10 mM sodium butyrate, and cells were incubated for an additional 16-24 h before use. Cells were growing at 37℃, 8% $CO_2$ and 100% humidity. Automated patch clamp recordings were conducted using the Syncropatch 384 PE platform (Nanion Technologies, Munich, Germany) at room temperature. We employed four-hole, 384-well recording chips with medium resistance ranging from 2 to 4 MΩ. Pulse generation and data collection was performed using the PatchController384 V.1.3.0 and DataController384 V1.2.1 software (Nanion Technologies). Whole-cell currents were filtered at 3 kHz and acquired at a sampling rate of 10 kHz. Access resistance and apparent membrane capacitance were estimated using the built-in protocols. Whole-cell currents were recorded from cells expressing TWIK-2 at room temperature with the internal solution containing 110 mM KF, 10 mM KCl, 10 mM NaCl, 10 mM HEPES, 10 mM EGTA, and pH adjusted to 7.2 with KOH. The external solution contained 44 mM NaCl, 100 mM KCl, 2 mM $CaCl_2$, 1 mM $MgCl_2$, and 5 mM glucose, buffered with either 10 mM HEPES (pH 7.4 adjusted with NaOH) or MES (pH 5.5 adjusted with NaOH). Voltage-dependent currents were measured by applying a holding potential of -90 mV. Outward currents of TWIK-2 were elicited using voltage steps ranging from -90 to +100 mV for a duration of 200 ms. Recordings were included from cells that exhibited a seal resistance greater than 100 MΩ, series resistance less than 10 MΩ, and stable cell capacitance throughout the recording session.

Time constants for activation of currents were determined and fit with a single exponential function using DataController384 (Nanion Technologies). The activation time constant (τ) was obtained by fitting the activation portion of the current waveforms between 10 and 90 % of the increment between holding (0 %) and peak values (100 %) in response to 200 ms voltage clamp steps. The $G/G_{MAX}$-$V$ curves were derived using the normalized chord conductance, calculated by dividing the maximum current elicited during a depolarizing step by the driving force derived from the calculated $K^+$ equilibrium potential. The $G/G_{MAX}$-$V$ curves were fitted with GraphPad Prism (v10.4.1) using a single Boltzmann function of the form: $y = 1/(1 + \exp[V_{1/2} - V]/k)$ where $y$ is the conductance normalized to the maximal conductance, $V_{1/2}$ is the half-activation potential, $V$ is the test voltage, and $k$ is the slope factor. For TWIK-2 concentration-response curves on the SyncroPatch 384PE, one compound concentration was applied to each well, and the concentration-response curve was calculated across the whole plate. For the analysis of the compound $IC_{50}$ value, concentration-response curves were fitted to a three-parameter sigmoid using GraphPad Prism (v10.4.1). For compound effect analysis, compound inhibition was calculated as the percentage of peak current (I) decrease from before compound application ($I_0$) to the end of 10 minutes of compound application ($I_{compound}$), both being normalized to the end of the experiment to the remnant currents after adding 10 mM $BaCl_2$ (full block). Statistical analyses were performed using the student's t test.

#### Drug and reagents

$K_{2P}$ modulator ML365 (Sigma-Aldrich SML2643), NPBA (MedChemExpress), ML335 (MedChemExpress) and BL1249 (Tocris) were diluted to 20 mM in sterile DMSO (Sigma-Aldrich) and stored in -80℃ before use.

**Figures S1 to S4**

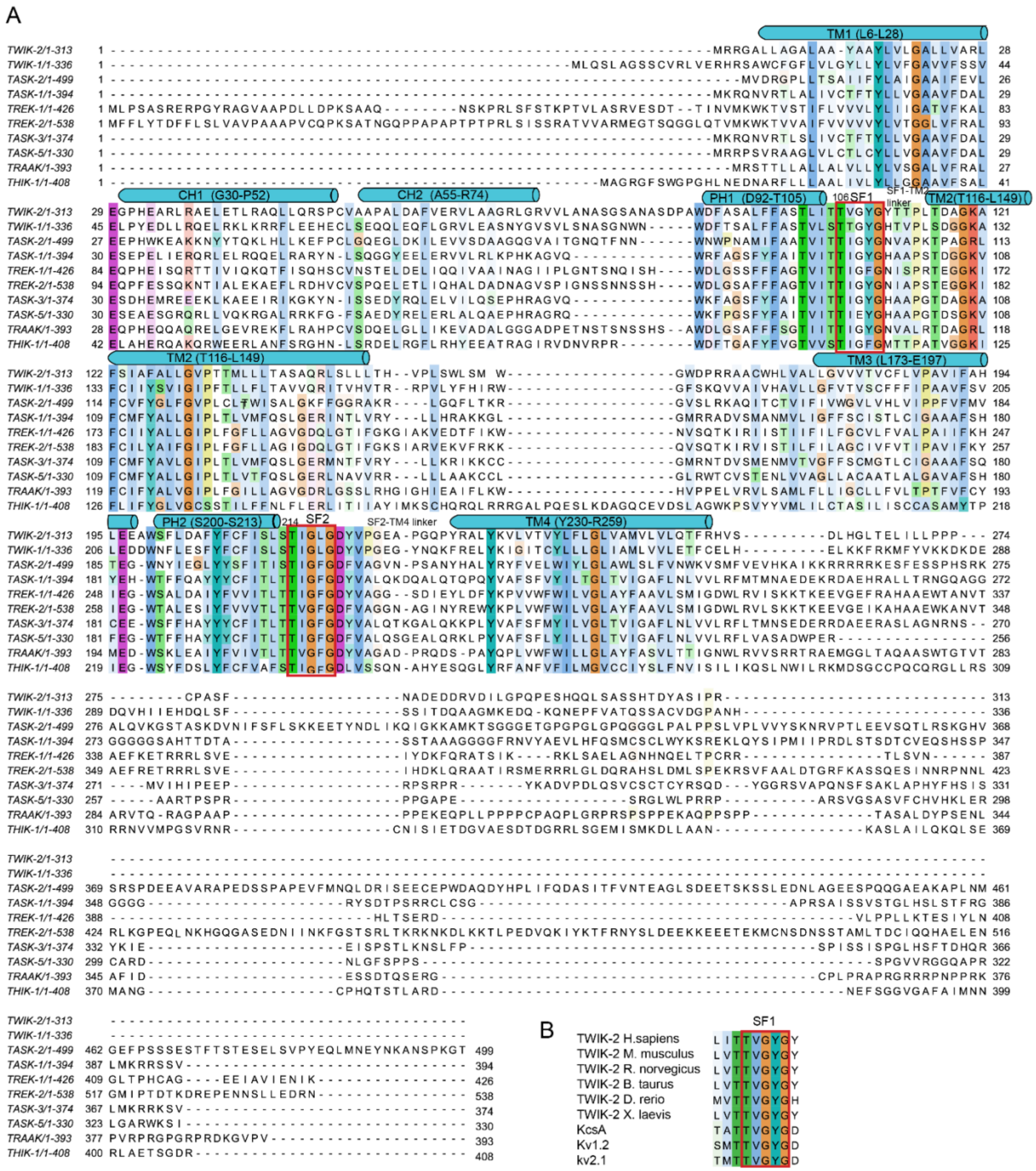

**Fig.S1. Sequence alignment of TWIK-2 with indicated channels. A**. Sequence alignment of TWIK-2 and other human K2P channels. Sequence alignments were colored by conservation in Jalview 2.11.4.1 (Procter, J. B. et al., 2021). TWIK-2 helical secondary structure elements are drawn above the sequences. Selectivity filters are boxed in red. Sequences are from human $K_{2P}$ channels: $K_{2P}$6.1 (TWIK-2), AAD22980.1,; $K_{2P}$1.1 (TWIK-1), AAB01688.1,; $K_{2P}$5.1 (TASK-2), AAC79458.1; $K_{2P}$3.1 (TASK-1), AAC51777.1; $K_{2P}$2.1 (TREK-1), AAD47569.1; $K_{2P}$10.1 (TREK-2), AAL95705.1; $K_{2P}$9.1 (TASK-3), AAF63708.1; $K_{2P}$15.1 (TASK-5), AAG33127.1; $K_{2P}$4.1 (TRAAK), AAF64062.1; $K_{2P}$13.1 (THIK-1), AAG32314.1. **B**. Sequence alignment of human TWIK-2 SF1 and other channels: TWIK-2 (*M.musculus* ) NM_001033525.3; TWIK-2 (*R.norvegicus*) NM_053806.2; TWIK-2 (*B.taurus*); NM_001205445.1; TWIK-2 (*D.rerio*) NM_001030074.2; TWIK-2 (*X.laevis*) NM_001096874.1;KcsA Z37969; Kv1.2 BC043564; Kv2.1 NM004975.

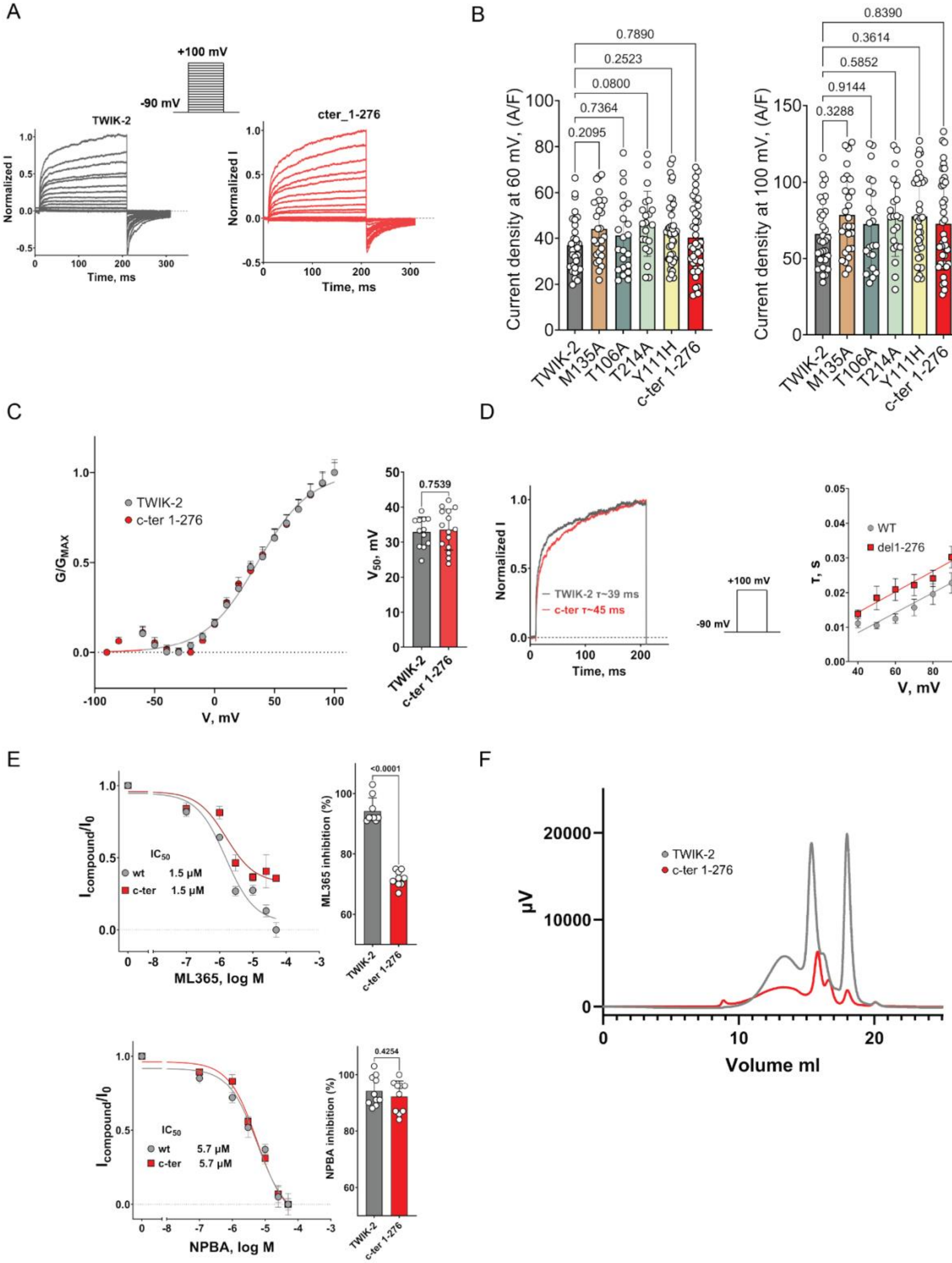
A
+100 mV
-90 mV
TWIK-2
cter_1-276
Normalized I
Time, ms
B
Current density at 60 mV, (A/F)
Current density at 100 mV, (A/F)
0.7890
0.2523
0.0800
0.7364
0.2095
0.8390
0.3614
0.5852
0.9144
0.3288
TWIK-2
M135A
T106A
T214A
Y111H
c-ter 1-276
C
TWIK-2
c-ter 1-276
G/G_MAX
V, mV
0.7539
V50, mV
D
Normalized I
TWIK-2 τ~39 ms
c-ter τ~45 ms
Time, ms
+100 mV
-90 mV
WT
del1-276
τ, s
V, mV
E
I_compound/I_0
IC50
wt 1.5 µM
c-ter 1.5 µM
ML365, log M
<0.0001
ML365 inhibition (%)
wt 5.7 µM
c-ter 5.7 µM
NPBA, log M
0.4254
NPBA inhibition (%)
F
TWIK-2
c-ter 1-276
µV
Volume ml

**Figure S2**. **Functional and expression analysis of ΔC-TWIK-2 (C-ter 1-276). A.** Families of voltage-gated current traces were recorded for TWIK-2 channels under high extracellular potassium ($K^+$) at pH 7.4. The normalized traces were obtained from 200-ms voltage steps ranging from -90 to +100 mV, starting from a holding potential of -90 mV using whole-cell patch clamp configuration**. B.** Scatter dot plots depict the current density (A/F) at +60 mV (top panel) and +100 mV (bottom panel) for TWIK-2 (n = 32) and ΔC-TWIK-2**,** n = 39). Data are expressed as mean ± SD. A one-way ANOVA with Dunnett's multiple comparisons was conducted to compare the wild-type channel against each mutant. **C**. The voltage-dependent activation curves for ΔC-TWIK-2, n = 15. The data are presented as mean ± SEM **D**. The half-maximal activation voltage shift (V50) for wild-type (n = 12) and ΔC-TWIK-2, n = 15 TWIK-2 channels is shown, with data expressed as mean ± SD. A one-way ANOVA with Dunnett's multiple comparisons was performed to evaluate differences between the wild-type and each mutant. **E**. Comparison of concentration-response curves of ML365 (top panel) and NPBA (bottom panel) on wild-type and ΔC-TWIK-2 channels. Data are presented as mean ± SEM: ML365 (n = 8) and NPBA (n = 7-8). Scatter dot plots display the current inhibition percentage at 50 µM for ML365 (n = 9) and NPBA (n = 10). The data are presented as mean ± SD. An unpaired t-test was conducted to compare the wild-type channel with ΔC-TWIK-2 channels. **F.** FSEC analysis of WT TWIK-2 and ΔC-TWIK-2 (C-ter 1-276) solubilized in 1% detergent using same amount of cell material.

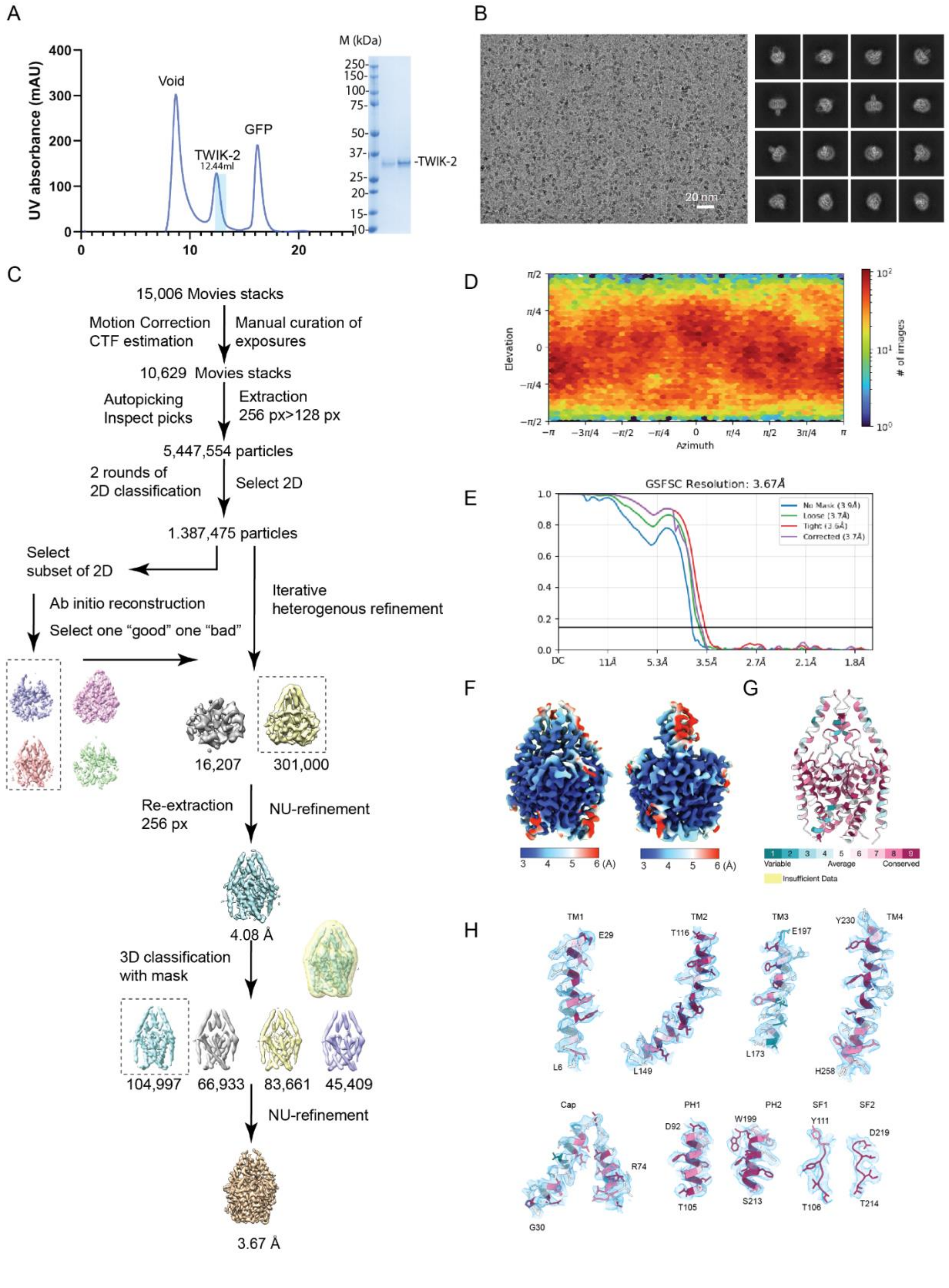
A
B
C
D
E
F
G
H
UV absorbance (mAU)
Void
TWIK-2
12.44ml
GFP
M (kDa)
-TWIK-2
20 nm
15,006 Movies stacks
Motion Correction
CTF estimation
Manual curation of
exposures
10,629 Movies stacks
Autopicking
Inspect picks
Extraction
256 px>128 px
5,447,554 particles
2 rounds of
2D classification
Select 2D
1.387,475 particles
Select
subset of 2D
Ab initio reconstruction
Select one "good" one "bad"
Iterative
heterogenous refinement
16,207
301,000
Re-extraction
256 px
NU-refinement
4.08 Å
3D classification
with mask
104,997
66,933
83,661
45,409
NU-refinement
3.67 Å
Elevation
Azimuth
### of images
GSFSC Resolution: 3.67Å
No Mask (3.9Å)
Loose (3.7Å)
Tight (3.6Å)
Corrected (3.7Å)
Variable
Average
Conserved
Insufficient Data
TM1
TM2
TM3
TM4
E29
L6
T116
L149
E197
L173
Y230
H258
Cap
R74
G30
PH1
D92
T105
PH2
W199
S213
SF1
Y111
T106
SF2
D219
T214

**Figure S3. Sample preparation and data processing of TWIK-2. A**. Size-exclusion chromatogram of TWIK-2 purified in DMNG, with peak eluted at 12.44 ml corresponding to dimer fraction of TWIK-2 shown on the SDS-PAGE. **B**. Representative raw micrograph and sample 2D class averages of TWIK-2. **C**. Data processing pipeline for TWIK-2 in DMNG. Details can be found in Methods. **D.** Particle distribution plot. E. Fourier shell correlation (FSC) curves for TWIK-2. **F**. TWIK-2 maps with estimated local resolution between 3-6 Å. **G**. Consurf[84] analysis of TWIK-2. **H**. Fit of the various secondary structural elements of TWIK-2 to the cryo-EM map. The cartoon is colored as per the evolutionary sequence conservation calculated by ConSurf[84]. The cryo-EM density is shown in light blue (display level 0.18 in ChimeraX).

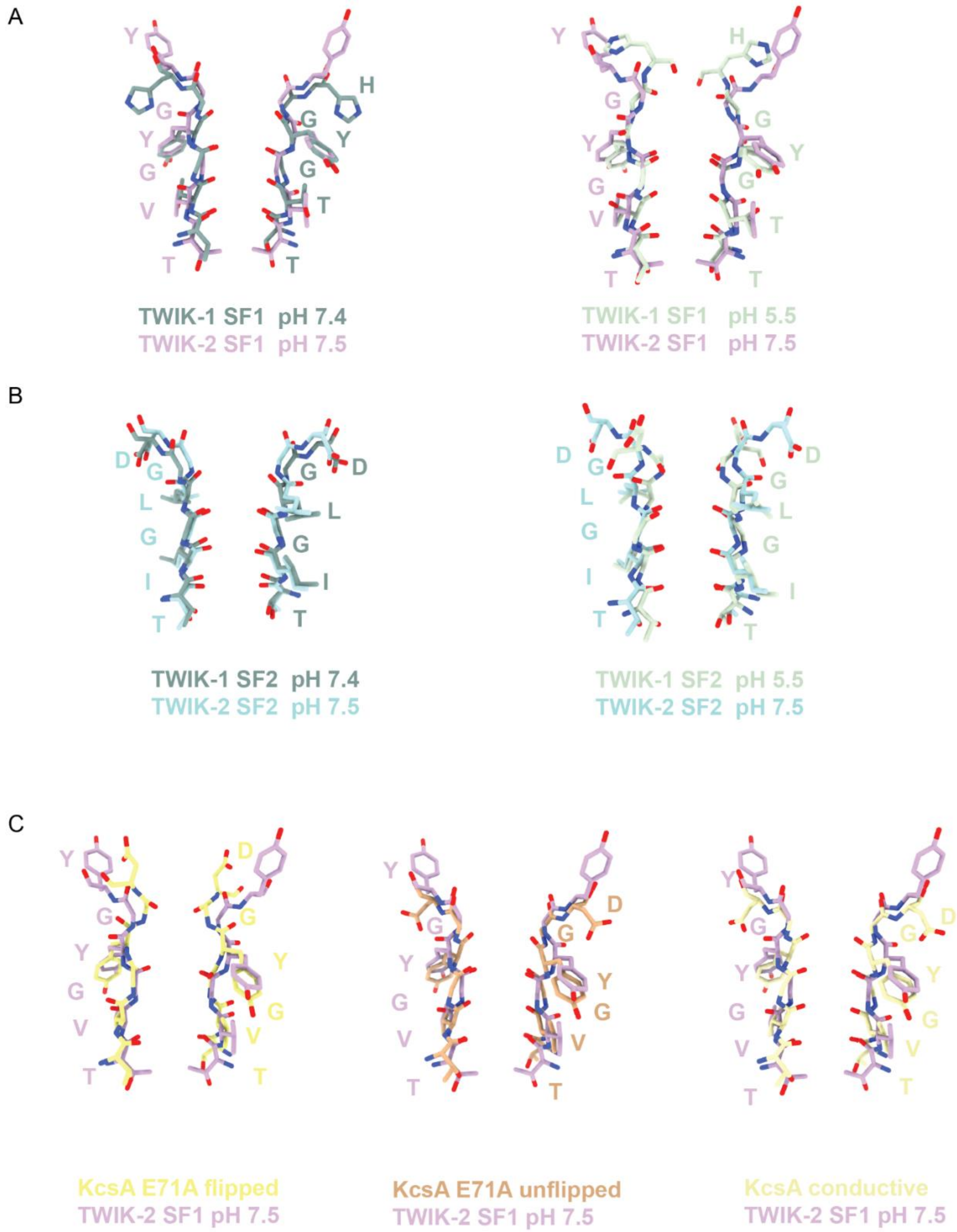


**Figure S4. Comparison of selectivity filters of TWIK-2 and other K$^+$ channels. A.** Overlaid TWIK-2 SF1 (floral) with TWIK-1(PDB: 7SK0) (left, moss) SF1 in high pH, open conformation and

TWIK-1 (PDB: 7SK1) (right, mint) SF1 in low pH, closed conformation **B.** Overlaid TWIK-2 SF2 (blue) with TWIK-1(PDB: 7SK0) (left, moss) SF2 in high pH, open conformation and TWIK-1 (PDB: 7SK1) (right, mint) SF2 in low pH, closed conformation. **C.** Overlaid TWIK-2 SF1 (Floral) with flipped KcsA E71A (PDB: 2ATK) (left, yellow), unflipped KscA E71A (PDB: 1ZW1) (middle, orange) and wild-type conductive KscA (PDB:1K4C) (right, light yellow).

**Table S1. Cryo-EM data collection, refinement, and validation statistics.**

| TWIK-2 (EMDB:47768) (PDB:9E94) | |
|---|---|
| **Data collection and processing** | |
| Magnification | 105,000 x |
| Voltage (kV) | 300 |
| Electron exposure (e–/Å2) | 60 |
| Defocus range (μm) | -0.8-2.5 |
| pixel size (Å) | 0.83 |
| Symmetry imposed | C1 |
| Initial particle images (no.) | 5,447,554 |
| Final particle images (no.) | 104,997 |
| Map resolution (Å) | 3.67 |
| FSC threshold | 0.143 |
| **Refinement** | |
| Model resolution (Å) | 3.67 |
| FSC threshold | 0.143 |
| Map sharpening B factor (Å2) | -135.9 |
| Model composition | |
| Nonhydrogen atoms | 3,381 |
| Protein residues | 440 |
| Potassium ion | 3 |
| D10 | 7 |
| *B* factors (Å2) | |
| Protein (min/max/mean) | 1.07/102.63/26.79 |
| ligands (min/max/mean） | 9.76/30.19/19.29 |
| R.m.s. deviations | |
| Bond lengths (Å) | 0.01 |
| Bond angles (°) | 1.563 |
| **Validation** | |
| MolProbity score | 1.75 |
| Clashscore | 6.18 |
| Poor rotamers (%) | 0 |
| Ramachandran plot | |
| Favored (%) | 94% |
| Allowed (%) | 6% |
| Disallowed (%) | 0 |

**References**


75. Goehring, A., Lee, C.-H., Wang, K.H., Michel, J.C., Claxton, D.P., Baconguis, I., Althoff, T., Fischer, S., Garcia, K.C., and Gouaux, E. (2014). Screening and large-scale expression of membrane proteins in mammalian cells for structural studies. Nat. Protoc. *9*, 2574–2585.

76. Punjani, A., Rubinstein, J.L., Fleet, D.J., and Brubaker, M.A. (2017). cryoSPARC: algorithms for rapid unsupervised cryo-EM structure determination. Nat. Methods *14*, 290–296.

77. Jamali, K., Käll, L., Zhang, R., Brown, A., Kimanius, D., and Scheres, S.H.W. (2023). Automated model building and protein identification in cryo-EM maps. Nature *631*, 610–616 (2024)

78. Emsley, P., Lohkamp, B., Scott, W.G., and Cowtan, K. (2010). Features and development of coot. Acta Crystallogr. D Biol. Crystallogr. *66*, 486–501.

79. Liebschner, D., Afonine, P.V., Baker, M.L., Bunkóczi, G., Chen, V.B., Croll, T.I., Hintze, B., Hung, L.W., Jain, S., McCoy, A.J., et al. (2019). Macromolecular structure determination using X-rays, neutrons and electrons: recent developments in Phenix. Acta Crystallogr. D Struct. Biol. *75*, 861–877.

80. Meng, E.C., Goddard, T.D., Pettersen, E.F., Couch, G.S., Pearson, Z.J., Morris, J.H., and Ferrin, T.E. (2023). UCSF ChimeraX: Tools for structure building and analysis. Protein Sci. *32*, e4792.

81. Smart, O.S., Neduvelil, J.G., Wang, X., Wallace, B.A., and Sansom, M.S. (1996). HOLE: a program for the analysis of the pore dimensions of ion channel structural models. J. Mol. Graph. *14*, 354–360, 376.

82. Schymkowitz, J., Borg, J., Stricher, F., Nys, R., Rousseau, F., and Serrano, L. (2005). The FoldX web server: an online force field. Nucleic Acids Res. *33*, W382–W388.

83. Laskowski, R.A., and Swindells, M.B. (2011). LigPlot+: multiple ligand-protein interaction diagrams for drug discovery. J. Chem. Inf. Model. *51*, 2778–2786.

84. Ashkenazy, H., Abadi, S., Martz, E., Chay, O., Mayrose, I., Pupko, T., and Ben-Tal, N. (2016). ConSurf 2016: an improved methodology to estimate and visualize evolutionary conservation in macromolecules. Nucleic Acids Res. 44, W344–W350.

85. Procter, J.B., Carstairs, G.M., Soares, B., Mourão, K., Ofoegbu, T.C., Barton, D., Lui, L., Menard, A., Sherstnev, N., Roldan-Martinez, D., et al. (2021). Alignment of biological sequences with Jalview. Methods Mol. Biol. 2231, 203–224.
